# Novel computational method for predicting polytherapy switching strategies to overcome tumor heterogeneity and evolution

**DOI:** 10.1101/086553

**Authors:** Vanessa D. Jonsson, Collin M. Blakely, Luping Lin, Saurabh Asthana, Victor Olivas, Matthew A. Gubens, Nikolai Matni, Boris C. Bastian, Barry S. Taylor, John C. Doyle, Trever G. Bivona

**Affiliations:** Department of Computing and Mathematical Sciences, California Institute of Technology, Pasadena, CA; Department of Medicine, University of California, San Francisco, CA; Helen Diller Family Comprehensive Cancer Center, University of California San Francisco, San Francisco, CA; Department of Pathology, University of California, San Francisco, CA; Human Oncology and Pathogenesis Program, Memorial Sloan Kettering Cancer Center; Department of Epidemiology and Biostatistics, Memorial Sloan Kettering Cancer Center; Marie-Josée and Henry R. Kravis Center for Molecular Oncology, Memorial Sloan Kettering Cancer Center, New York, NY; Department of Control and Dynamical Systems, California Institute of Technology, Pasadena, CA; Department of Electrical Engineering, California Institute of Technology, Pasadena, CA; Department of Biological Engineering, California Institute of Technology, Pasadena, CA; Cellular and Molecular Pharmacology, University of California, San Francisco, CA

## Abstract

The success of targeted cancer therapy is limited by drug resistance that can result from tumor genetic heterogeneity. The current approach to address resistance typically involves initiating a new treatment after clinical/radiographic disease progression, ultimately resulting in futility in most patients. Towards a potential alternative solution, we developed a novel computational framework that uses human cancer profiling data to systematically identify dynamic, pre-emptive, and sometimes non-intuitive treatment strategies that can better control tumors in real-time. By studying lung adenocarcinoma clinical specimens and preclinical models, our computational analyses revealed that the best anti-cancer strategies addressed existing resistant subpopulations as they emerged dynamically during treatment. In some cases, the best computed treatment strategy used unconventional therapy switching while the bulk tumor was responding, a prediction we confirmed in vitro. The new framework presented here could guide the principled implementation of dynamic molecular monitoring and treatment strategies to improve cancer control.

## 2 Introduction

Targeted cancer therapies are effective for the treatment of certain oncogene-driven solid tumors, including non-small cell lung cancers (NSCLCs) with activating genetic alterations in EGFR (epidermal growth factor receptor), ALK (anaplastic lymphoma kinase), BRAF, and ROS1 kinases [1, 2, 3]. However, inevitably resistance to current targeted therapies emerges, typically within months of initiating treatment and remains an obstacle to long-term patient survival [1, 2, 3, 4]. The presence and evolution of tumor genetic heterogeneity potentially underlies resistance and also limits the response to successive therapeutic regimens that are used clinically in an attempt to overcome resistance in the tumor after it has emerged [4, 5, 6, 7]. Indeed, while a targeted therapy may be effective in suppressing one genomic subclone within the tumor, other clones may be less sensitive to the effects of the drug. Thus, through selective pressures, resistant populations can emerge and promote tumor progression. Moreover, the current paradigm of solid tumor treatment is largely based on designing fixed (static) treatment regimens that are deployed sequentially as either initial therapy or after the clear emergence of drug-resistant disease, detected by clinical and radiographic measures of tumor progression. In contrast, designing dynamic treatment strategies that switch between targeted agents (or combinations thereof) in real time in order to suppress the outgrowth of rare or emergent drug-resistant subclones may be a more effective strategy to continually suppress tumor growth and extend the duration of clinical response. Thus, there is a need to identify principled approaches for the predictive design of effective combination (poly)therapy strategies to pre-empt the growth of multiple tumor subclones actively during treatment.

Mathematical modeling, analysis and computational simulations of tumor growth, heterogeneity and inhibition by various therapeutic modalities has long been employed as a method to provide insight into evolutionary outcomes and effective treatment strategies. Such modeling may include the use of stochastic [8, 9, 10] or deterministic differential equation implementations [11, 12] to propose static or sequential treatment strategies that delay resistance in various cancer models. Recent studies by Zhao et al. [13, 14] incorporate the use of mathematical optimization, a fundamental subject in engineering design to predict static combination therapies that effectively address heterogeneity in a lymphoma model. Complementary engineering techniques from optimal control theory provide an additional theoretical framework to design dynamic drug scheduling regimens in the context of dynamical systems models of cancer heterogeneity and evolution. The application of optimal control theory to treatment design has a history dating back to the 1970s [15, 16] with more recent examples including that of scheduling angiogenic and chemotherapeutic agents [17] or immuno- and chemotherapy combinations [18]. While mathematical modeling and engineering methods have been used extensively to inform treatment strategy design, a significant drawback to prior work in the field is that the underlying computational framework(s) have not conjointly accomplished the following important aims: (1) allowing for the systematic principled design of dynamic treatment strategies using experimentally identified models of tumor dynamic behaviors; and (2) developing quantitative methods that allow for the exploration of the robustness of predicted treatment strategies with respect to multiple common challenges in real-world patients, such as tumor heterogeneity and fluctuations in drug concentrations.

Here, we present a novel approach that combines a mathematical model of the evolution of tumor cell populations with parameters identified from our experimental data and an engineering framework for the systematic design of polytherapy scheduling directed at the following unresolved issues in the field: (1) how tumor genetic composition and drug dose constraints affect the long term efficacy of combination strategies, (2) how optimal scheduling of combination small molecule inhibitors can help to overcome heterogeneity, genomic evolution and drug dose fluctuations, and (3) how serial tumor biopsy or blood-based tumor profiling scheduling in patients can be timed appropriately. To tackle these questions, we developed an integrated experimental and computational framework that solves for candidate combination treatment strategies and their scheduling given an initial polyclonal tumor and allows the exploration of treatment design trade offs such as dosage constraints and robustness to small fluctuations in drug concentrations. This methodology is rooted in optimal control theory and incorporates an experimentally derived mathematical model of evolutionary dynamics of cancer growth, mutation and small molecule inhibitor pharmacodynamics to solve for optimal drug scheduling strategies that address tumor heterogeneity and constrain drug-resistant tumor evolution. Our key new insights include (1) heterogeneous tumor cell populations are better controlled with switching strategies; indeed, static two-drug strategies are unable to effectively control all tumor subpopulations in our study; (2) constant combination drug strategies are less robust to perturbations in drug concentrations for heterogeneous tumor cell populations, and hence more likely to lead to tumor progression; (3) countering the outgrowth of subclonal tumor populations by switching polytherapies even during a bulk tumor response can offer better tumor cell population control, offering a non-intuitive clinical strategy that pro-actively addresses molecular progression before evidence of clinical or radiographic progression appears.

## 3 Results

### 3.1 The presence and evolution of intratumoral genetic heterogeneity in a patient with EGFR-mutant lung adenocarcinoma

To explore the utility of our approach, we focused on EGFR-mutant lung adenocarcinoma. Many mechanisms of resistance to EGFR-targeted therapies in lung adenocarcinoma are well characterized [19]. Furthermore, tumor heterogeneity and multiple resistance mechanisms arising in a single patient can occur [2, 19]. Thus, overcoming polygenic resistance is of paramount importance in this disease and will likely require a non-standard approach. To illustrate this point, we investigated the molecular basis of targeted therapy resistance in a 41-year old male never-smoker with advanced EGFR-mutant (L858R) lung adenocarcinoma. This patient responded to first-line treatment with erlotinib but progressed on this therapy within only four months after initial treatment, instead of the typical 9-12 month progression free survival observed in EGFR-mutant lung adenocarcinoma patients. We reasoned that genomic analysis of this patient’s outlier clinical phenotype could reveal the molecular pathogenesis of suboptimal erlotinib response. Using a custom-capture assay [20, 21], we deeply sequenced the coding exons and selected introns of 389 cancer-relevant genes in both the pre-treatment and the erlotinib-resistant tumor specimen and matched normal blood to identify somatic alterations that could mediate resistance (Materials and Methods). Exome sequencing of the pre-treatment specimen confirmed the presence of the EGFR^L858R^ mutant allele that was identified through prior clinical PCR-based sequencing of this EGFR^L858R^ specimen (data not shown), and additionally revealed mutant allele-specific focal amplification of the EGFR coding locus that resulted in a high allelic frequency (95% variant frequency) (Fig. 1B-C). We discovered a rare concurrent subclone in the treatment-naïve tumor with a BRAF^V600E^ mutation (6% variant frequency; Fig. 1B). This observation is consistent with a recent report of a BRAF^V600E^ mutation in an erlotinib-resistant lung adenocarcinoma specimen [22] and recent data indicating that EGFR-mutant lung adenocarcinoma cells can often develop EGFR TKI resistance through RAF-MEK-ERK pathway activation [23]. The frequency of the subclonal BRAF^V600E^ mutation increased approximately 10-fold upon acquired erlotinib resistance, from 6% to 60% in the primary and recurrent tumor, respectively (Figure 1C). This increase in the BRAF^V600E^ allelic fraction was likely due to the expansion of the BRAF^V600E^ subclone, given that we found no evidence that this increased frequency occurred as a result of focal BRAF amplification in the resistant tumor (Figure 1C). Beyond the outgrowth of mutant BRAF, we identified two additional genetic alterations in the resistant tumor that could contribute to EGFR TKI resistance: focal amplification of 7q31.2 encoding MET in the resistant tumor cells, a low frequency EGFR^T790M^ mutation (14% variant frequency) (Fig. 1B-C). All candidate somatic mutations and focal copy number amplifications conferring resistance to erlotinib therapy (in EGFR, BRAF, and MET) were confirmed by independent, validated DNA sequencing and FISH assays (data not shown). Thus, erlotinib therapy acted as a selective pressure for the evolution of multiple concurrent clonal and subclonal genetic alterations that could cooperate to drive rapid drug-resistant disease progression in EGFR-mutant lung adenocarcinoma.

**Figure 1:**
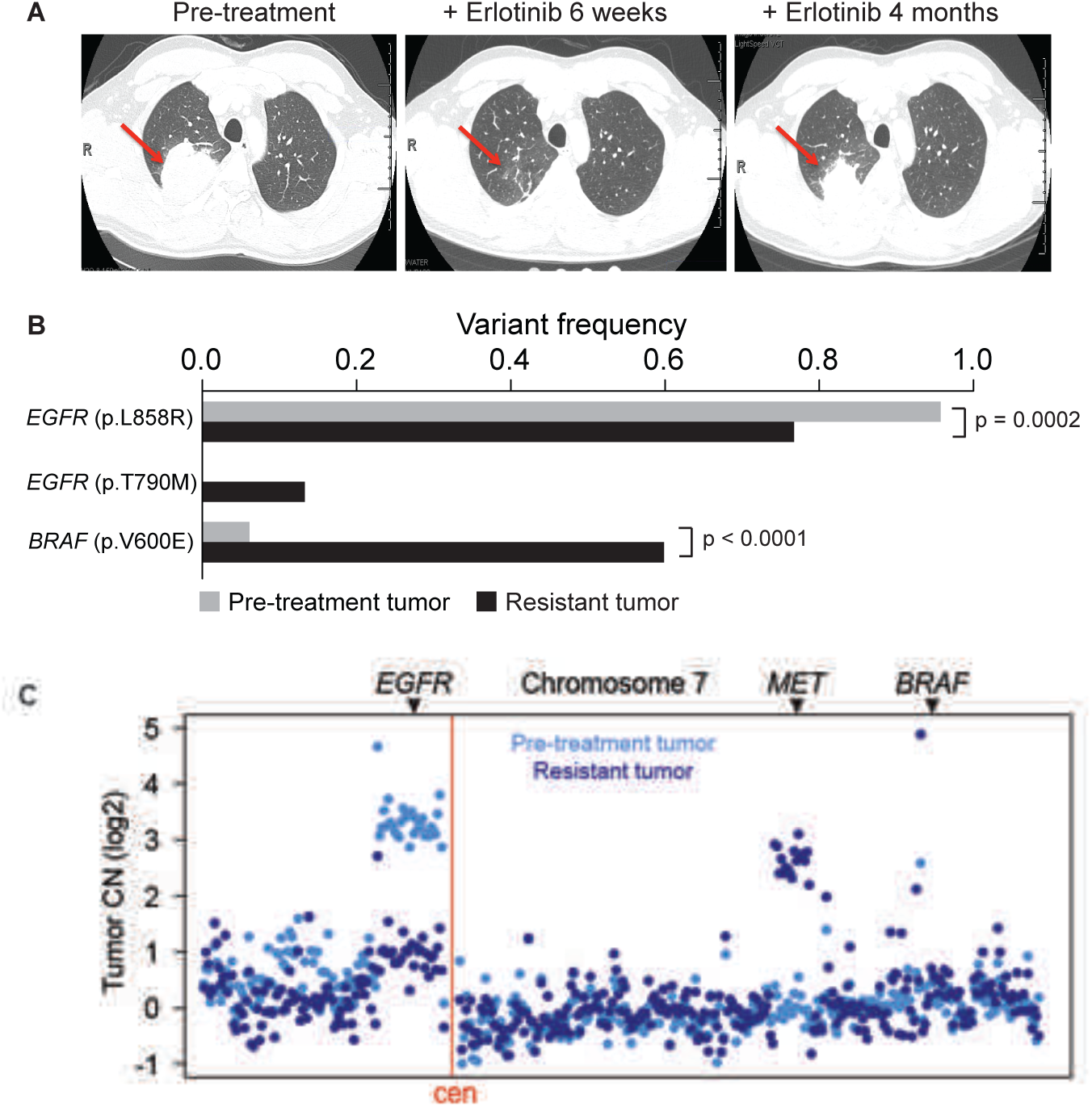
Concurrent genetic alterations drive rapid resistance to EGFR TKI treatment in EGFR-mutant lung adenocarcinoma. (A) Computed tomography indicates the clinical course and timeline of disease in the patient with rapid progression on EGFR TKI therapy and shows the EGFR-mutant lung adenocarcinoma (red arrows) analyzed both prior to erlotinib treatment and upon resistance at 4 months. (B) Key somatic mutations identified by exon-capture and deep sequencing of the pre- and post-treatment tumor in (A) demonstrating concurrent alterations in EGFR and BRAF and the frequency of each mutation in pre- and post- treatment tumor samples. P-values indicated as determined by a two-tailed Fischer’s exact test. (C) DNA copy number alterations inferred from exon-capture and sequencing data indicate the focal amplification of the EGFR^L858R^-mutant allele was lost upon acquired resistance while the patient’s resistant tumor gained a focal amplification of MET, with no change in BRAF (relative positions indicated, chromosome 7).

### 3.2 Analysis of clonal concurrence and resistance

While BRAF^V600E^, MET activation, and EGFR^T790M^ can individually promote EGFR TKI resistance [22, 24, 25], the therapeutic impact of the concurrence of these alterations we uncovered has not been characterized. Therefore, we studied the effects of BRAF^V600E^, MET activation, and EGFR^T790M^, alone or in combination, on growth and therapeutic response in human EGFR-mutant lung adenocarcinoma cellular models. First, we found that expression of V600E but not wild-type (WT) BRAF promoted resistance to erlotinib in 11-18 cells that endogenously express EGFR^L858R^ (Fig. S1). This erlotinib resistance in BRAF^V600E^-expressing EGFR-mutant 11-18 cells was overcome by concurrent treatment with erlotinib and selective inhibitors of either BRAF or MEK (vemurafenib [26] and trametinib [27] respectively (Fig. S2, S3). We next used the 11-18 system to test the effects of MET activation by hepatocyte growth factor (HGF), which phenocopies the effects of MET amplification in EGFR TKI resistance[25, 28] on therapeutic sensitivity. We found that MET activation not only promoted erlotinib resistance in parental 11-18 cells but also enhanced the effects of BRAF^V600E^ expression on erlotinib resistance in these cells (Fig. S1). This resistance induced by MET activation in 11-18 parental and BRAF^V600E^-expressing cells was accompanied by increased phosphorylation of MEK, ERK, and AKT (Fig. S3). Treatment with the MEK inhibitor trametinib, but not the BRAF inhibitor vemurafenib or the MET inhibitor crizotinib, overcame erlotinib resistance and inhibited phospho-ERK in MET-activated BRAF^V600E^-expressing 11-18 cells (Fig. S3), providing a rationale for polytherapy against EGFR and MEK in EGFR-mutant tumors with activating co-alterations in MET and BRAF.

Given that we found a rare EGFR^T790M^ subclone in the polyclonal resistant tumor, we next explored whether BRAF^V600E^ expression could promote resistance to EGFR TKI treatment in H1975 human lung adenocarcinoma cells that endogenously express EGFR^T790M^ and EGFR^L858R^. We observed that BRAF^V600E^ modestly decreased sensitivity to afatinib, an approved irreversible EGFR kinase inhibitor effective against EGFR^T790M^ [29], and that this effect of BRAF^V600E^ on afatinib sensitivity was blunted by vemurafenib (Fig. S4). Together, our data indicate that erlotinib therapy induced the evolution of multiple concurrent events that re-shaped the polyclonal tumor genetic landscape during the onset of resistance; resistance could be overcome by polytherapy against both EGFR and MAPK signaling in preclinical models.

### 3.3 Polytherapy Provides Temporary Response in Heterogeneous or MET Activated Tumors

While we conducted a finite set of experiments to test various rational drug combinations that could address the heterogeneous basis of resistance in this patient’s disease, this approach is not easily scaled; further, it is not readily feasible to explore all possible drug combinations and drug doses over a continuous range or anticipate the effects of the myriad of possible tumor subcompositions on tumor control under treatment using cell-based assays alone. Therefore, we sought to provide a more general and scalable framework for understanding the impact of each genetically-informed targeted therapy strategy on the temporal evolution of the multiple concurrent EGFR-mutant tumor cell subclones present in this patient, as a potentially more generalizable platform. We developed an ordinary differential equation (ODE) model of tumor growth, mutation and selection by small molecule inhibitors with parameters identified from experimental data (Fig. 2A-B and Equation S1) and interrogated it to uncover the limitations of the targeted treatments in the context of tumor heterogeneity and evolution. We first confirmed that our model was able to capture the essential tumor population dynamics by showing a qualitative equivalence between the patient’s clinical course and our model simulation of similar tumor subpopulations consisting of 94 % EGFR^L858R^, 6% BRAF^V600E^ and assuming the existence of a very low initial frequency of 0.01% MET amplification of EGFR^L858R^, BRAF^V600E^ and EGFR^T790M^ in the presence of 1 *µ*M erlotinib (Fig. 3A-B).

**Figure 2:**
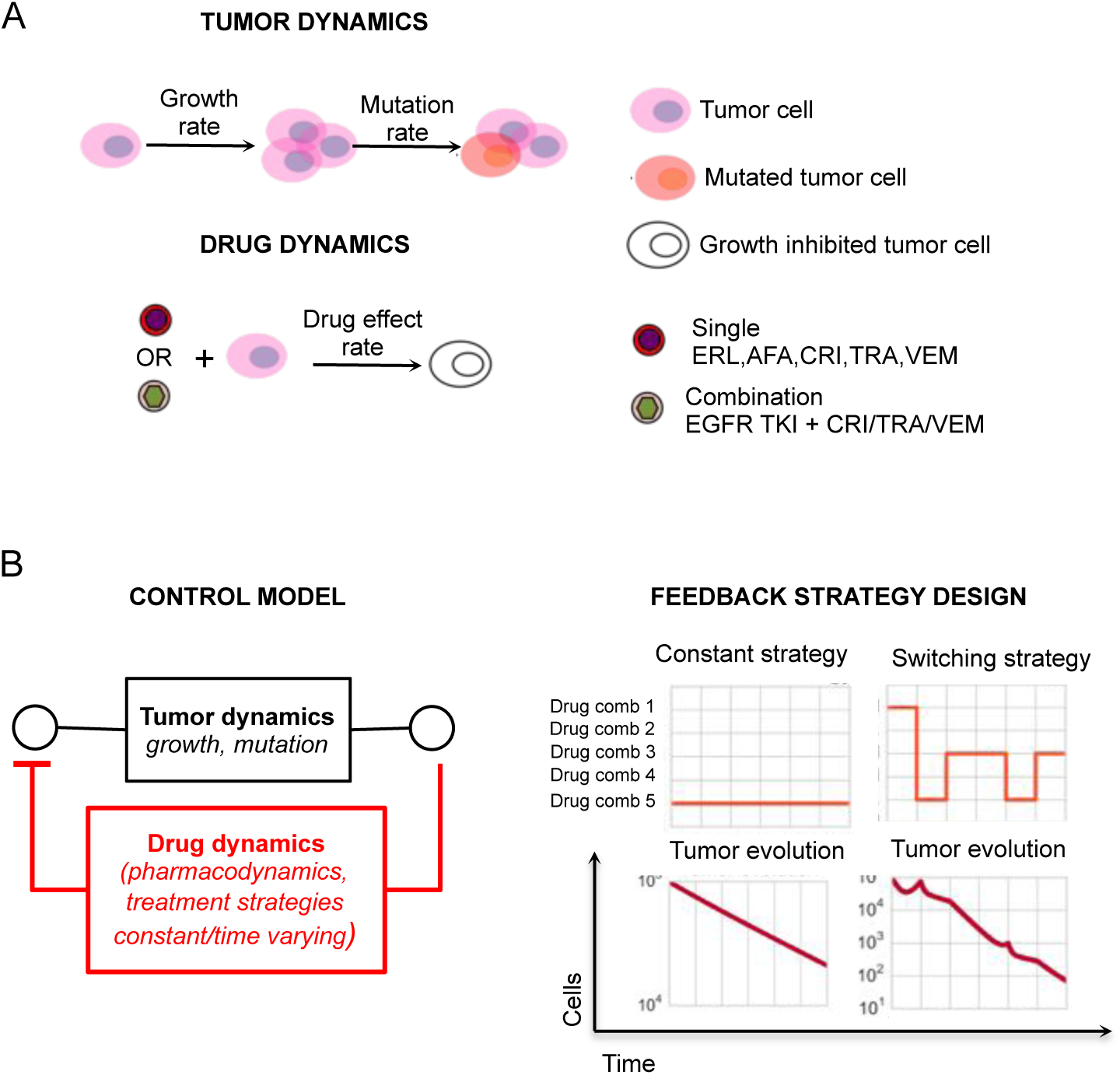
Designing treatment strategies to control tumor cell dynamics. (A) A depiction of the growth, mutation and drug effect model representing the evolutionary dynamics of lung adenocarcinoma in the presence of small molecule inhibitors, erlotinib (ERL), afatinib (AFA), crizotinib (CRI), trametinib (TRA) and vemurafenib (VEM). The corresponding ordinary differential equation model (ODE) is specified in mathematical detail in the Supplementary Information, Equation (S1). Drug effect curves were determined for 11-18 and H1975 cell lines specified for both single drugs and combinations of varying concentrations of one EGFR TKI (erlotinib or afatinib), with fixed concentrations of either 5 *µ*M vemurafenib, 0.5 *µ*M trametinib or 0.5 *µ*M crizotinib (SI, Fig. S1-S4). (B) The design of constant or switching feedback strategies to control the dynamics of lung adenocarcinoma is approached as an optimal control problem. The treatment strategy design algorithm (SI, Section 2) solves for feedback strategies that minimize tumor cell growth over the course of the treatment.

**Figure 3:**
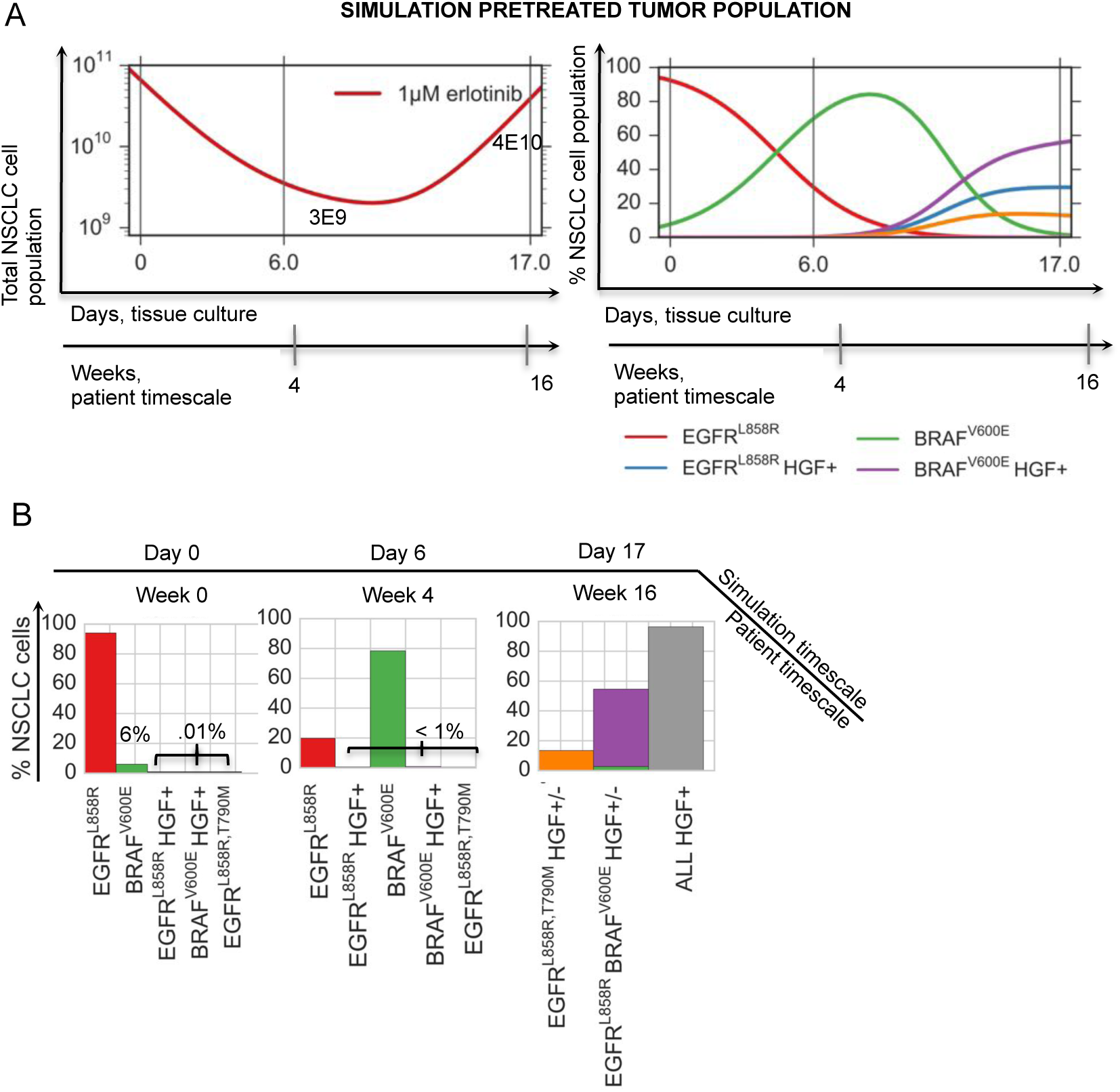
Mathematical simulation qualitatively captures the patient’s evolution on erlotinib. (A) A simulation of the mathematical model of lung adenocarcinoma evolution (SI, Equation (S1)) in the presence of 1 *µ*M erlotinib, given the patient-derived pretreatment initial tumor cell subpopulations (94 % EGFR^L858R^, 6% BRAF^V600E^, 0.01% MET amplification of EGFR^L858R^, BRAF^V600E^ and EGFR^T790M^). (B) Tumor cell populations present at day 0, 6 and 17 of the simulation in (A), including the total HGF+ cell population at day 17 (gray). The model qualitatively captures a possible evolutionary trajectory and results in a similar final tumor cell composition as that of the patient, (B) day 17 vs. Figure 1, (B) and (C).

To systematically explore the utility of many different drug combination regimens to overcome polygenic resistance, we used our computational model to calculate the efficacy of clinically relevant doses of erlotinib and afatinib in combination with either crizotinib, trametinib or vemurafenib on the growth of parental 11-18 and H1975 cells EGFR mutant cell lines. We found that most polytherapies could address only certain subpopulations (Fig. 4A). For example, the afatnib/trametinib combination elicited a complete response for a representative heterogeneous MET- tumor cell population comprised of (89% EGFR^L858R^, 10% EGFR^L858R^ BRAF^V600E^, 1% EGFR^L858R, T790M^) compared to rapid progression for its MET activated analog (Fig. 4C). Moreover, we computed the concentrations of erlotinib or afatinib in combination that could guarantee a progression-free response for both MET activated or MET neutral tumor cell populations (SI, Mathematical Methods) and found that in many cases, the concentrations were considerably higher than clinically feasible (due to either known pharmacokinetic limitations or dose limiting toxicities) (Fig. 4B).

**Figure 4:**
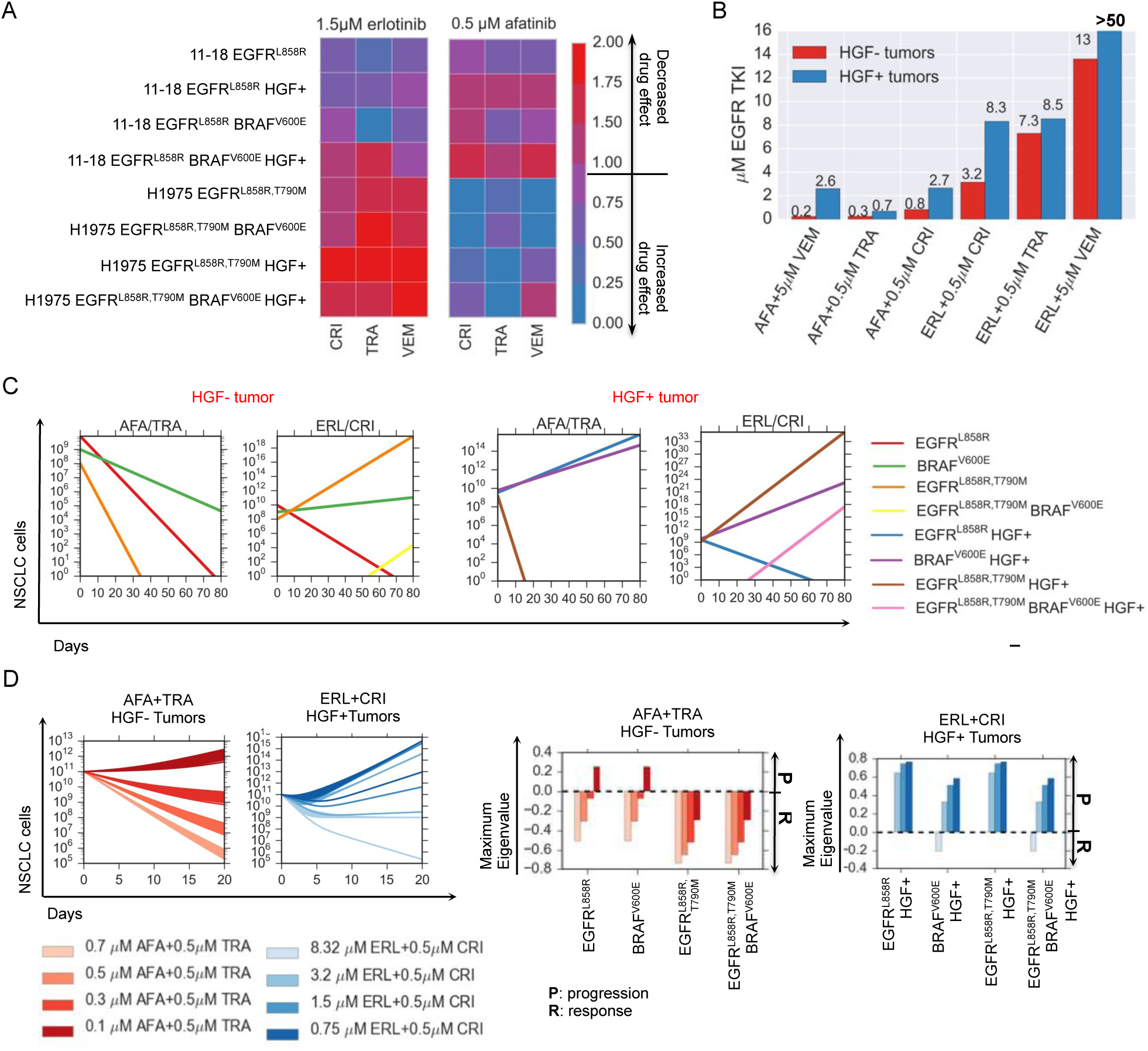
Modeling pharmacodyamic effects of concurrent BRAF^V600E^ expression and MET activation in EGFR-mutant lung adenocarcinoma cells and their implication on progression. (A) Drug efficacy as measured by the effect of 1.5 *µ*M erlotinib or 0.5 *µ*M afatinib in combination with either 0.5 *µ*M MET inhibitor crizotinib, 0.5 *µ*M MEK inhibitor trametinib or 5 *µ*M BRAF inhibitor vemurafenib on cell growth (SI, Equation S1) of parental 11-18 EGFR^L858R^-positive lung adenocarcinoma cells or those cells engineered to express mutations listed above and treated with 0 or 50 ng/ml HGF. (B) Concentrations of EGFR TKIs afatinib and erlotinib in combination with either 0.5 *µ*M crizotinib, 0.5 *µ*M trametinib or 5 *µ*M vemurafenib that guarantee progression free tumor reduction for any HGF- or HGF+ initial tumor subpopulations according to the model, measured by the minimum concentration of erlotinib or afatinib that results in exponential stability of the evolutionary dyanmics model (SI, Section 3.2). (C) Simulations of the lung adenocarcinoma model for combinations of 0.5 *µ*M afatinib+0.5 *µ*M trametinib and 1.5 *µ*M erlotinib+0.5 *µ*M crizotinib for the HGF- and HGF+ tumors specified. (D) (Left) Simulations of the evolutionary dynamics of different HGF- lung adenocarcinoma initial tumor subpopulations with a constant treatment of 0.7 *µ*M, 0.5, 0.3 or 0.1 *µ*M afatinib in combination with 0.5 *µ*M of trametinib (red) and of different HGF+ lung adenocarcinoma initial tumor subpopulations with a constant treatment of 8.32 *µ*M, 3.2 *µ*M, 1.5 *µ*M or 0.75 *µ*M erlotinib in combination with 0.5 *µ*M crizotinib (blue). (Right) Maximum eigenvalue decompositions (SI, Section 3.2) classify which subpopulationss can lead to progression at different concentrations of EGFR TKI for the afatinib+trametinib combination and the erlotinib+crizotinib combination.

To better understand the efficacy of the combination therapy over time, we sought to classify which initial tumor cell subpopulations could eventually lead to therapeutic failure when treated with different concentrations of EGFR TKIs in combination with crizotinib, trametinib or vemurafenib. We defined the evolutionary stability of a subpopulation as the worst-case evolutionary outcome, in each case where the particular subpopulation is present upon treatment initiation. More precisely, the evolutionary stability is the maximum eigenvalue of each evolutionary branch downstream of the subclone (SI, Section 3.2). This approach provides an assessment of which subclones present in the initial tumor cell population are likely to lead to overall progression (a positive evolutionary stability) versus those that lead to response (a negative evolutionary stability) when treated with a particular combination therapy. Our analysis confirms that progression-free response on combination therapies is sensitive to both EGFR TKI concentration and dependent on whether pre-existent subpopulations are effectively targeted at these concentrations (Fig. 4D and Fig. S5-S7). Overall, this analysis revealed that combinations of two signal transduction inhibitors had limited effectiveness in durably controlling resistance over a longer time horizon.

### 3.4 Engineering Drug Scheduling to Control Tumor Evolution

We next explored how the rational design of combination drug scheduling strategies could address this issue. Experimental studies have recently proposed drug pulsing [30] or drug switching [10] as a strategy to delay the growth of certain cancers. To this end, we proposed a novel methodology rooted in engineering principles to design drug scheduling strategies that best control the growth and evolution of tumor cell populations. In particular, we apply concepts from optimal and receding horizon control theory to our experimentally integrated model of lung adenocarcinoma evolution to compute treatment strategies that minimize tumor cell populations over time. Our algorithm allows for the specification of treatment design constraints such as maximum dose, the time horizon over which the treatment strategy is applied and the switching horizon, that is the minimum time over which one particular treatment can be applied. This algorithm can be extended to include other drug related characteristics and treatment design constraints. In addition, the framework allows for the analysis of tradeoffs between these aspects of the design space as well as others, such as how robust the predicted treatment strategies are with respect to uncertainties in the model or perturbations in drug dosages.

For a predetermined time and minimum switching horizon, we define an optimal control problem (SI, Algorithm 1) and solve for the drug combination that best minimizes the existing tumor cell subpopulations for every receding switching horizon. Given that any one polytherapy is unlikely to be simultaneously effective against all subpopulations, the resulting optimal strategy, which maximizes the response of the tumor cells present at every time horizon (SI, Mathematical Methods), is potentially one that switches between drug combinations, at defined time points during the treatment course.

As proof-of-principle, we determined which drug scheduling regimens could maximally reduce different initial tumor cell populations by solving our control problem for different allowable switching horizons over a thirty day period (Fig. 5). The afatinib/trametinib combination was the optimal constant strategy for tumor cell populations harboring the EGFR^L858R,T790M^ mutation, and although this strategy invoked progression free response in HGF- tumor cell populations, most L858R HGF+ tumor cell populations progressed on the therapy over thirty days (Fig. 5A vs 5C and Fig. S6AB). For the HGF- tumor population comprised of 89% EGFR^L858R^, 10% EGFR^L858R^BRAF^V600E^ and 1% EGFR^L858R,T790M^, the optimal constant strategy provided overall response leaving a dominant EGFR^L858R^BRAF^V600E^ tumor subpopulation present whereas the optimal ten day switching strategy provided an enhanced response over the constant strategy by alternately targeting EGFR^L858R^ and EGFR^L858R, T790M^ tumor cell subpopulations (Fig. 5B). In the case of the HGF treated tumor cell distribution consisting of 90% EGFR^L858R^ and 10% EGFR^L858R,T790M^, a constant combination of afatinib/trametinib was effective against the EGFR^L858R,T790M^, HGF+ subpopulation despite overall progression due to the outgrowth of the EGFR^L858R^, HGF+ tumor cell population, whereas a five day switching regimen between afatinib/trametinib and erlotinib/crizotinib combinations alternately targeted the HGF+ EGFR^L858R,T790M^ and the EGFR^L858R^ subpopulations (Fig. 5B and Fig. 3A) leading to overall response.

**Figure 5:**
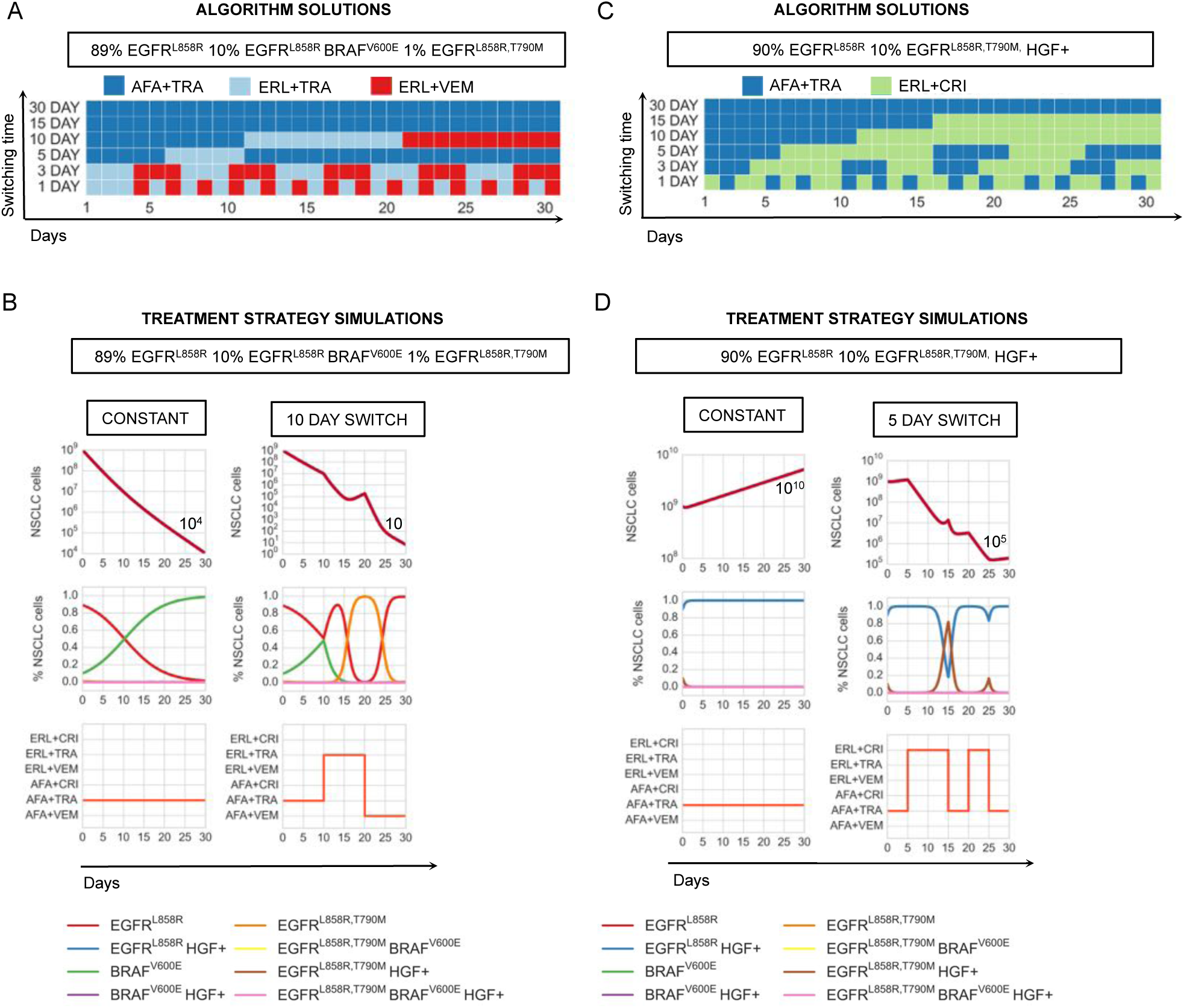
Optimal drug scheduling strategies solved by Algorithm 1 (SI, Section 2.2) for representative initial tumor cell distributions (A),(C), for a 30 day timeframe and 30, 15, 10, 5, 3 and 1 day minimum switching horizons, give one EGFR TKI, either 1.5 *µ*M erlotinib (ERL) or 0.5 *µ*M afatinib (AFA) in combination with either 5 *µ*M vemurafenib (VEM), 0.5 *µ*M trametinib (TRA) or 0.5 *µ*M crizotinib (CRI) and corresponding simulations (B),(D) of the lung adenocarcinoma evolutionary dynamics for a subset of optimal drug scheduling strategies.

More generally, the optimal constant strategies determined by our algorithm are combinations that best minimize existing tumor cell subpopulations at every switching horizon. In particular, a greater reduction in tumor cells can be achieved by switching between therapies that alternately target different subpopulations, even while there is overall response in the tumor (Fig. 5A). This finding suggests a non-intuitive approach to the clinical management of solid tumors that would represent a departure from the current standard clinical practice. Our model suggests an advantage to switching treatments pro-actively even during a bulk tumor response, while the current paradigm in the field is to switch from the initial treatment to a new drug(s) only after there is clear evidence of radiographic or clinical progression on the initial treatment.

To understand the potential benefits of switching strategies in tumors with different initial genetic heterogeneity, we computed the optimal switching strategies for a subset of tumor cell distributions and compared them to their corresponding computed optimal constant strategies. We found that the larger the number of subclones present in the initial tumor, the more beneficial even a small number of switches could be for overall tumor cell population control (Fig. 6A and Fig. S8A). For a highly heterogeneous tumor cell population comprised of HGF treated 89% EGFR^L858R^, 10% EGFR^L858R^BRAF^V600E^, 1% EGFR^L858R,T790M^ mutations, the predicted fifteen day switching therapy (afatinib/trametib followed by erlotinib/crizotinib) provides an immediate benefit versus the predicted constant treatment strategy (afatinib/trametinib), yielding a 10-fold decrease in final tumor population. By contrast, for a more homogeneous tumor consisting of 90% EGFR^L858R^, 10% EGFR^L858R,T790M^, the optimal predicted 30, 15 and 10 day switching strategies are indistinguishable from the constant therapy strategy for population control. Our predictions indicate that a similar 10-fold reduction in final population (similar to that achieved in the heterogeneous tumor instance analyzed above) is achieved only with a more rapid, five day switching strategy for this more homogeneous tumor population (afatinib/trametinib, then alternating between erlotinib/trametinib and afatinib/vemurafenib). These findings emphasize our results that while polytherapy may a provide response in some subsets of tumor cell populations, it provides only a temporary or no response in heterogeneous or MET activated tumors; in these cases, even minimal therapy switching can provide an immediate and more substantial benefit for overall tumor population control.

**Figure 6:**
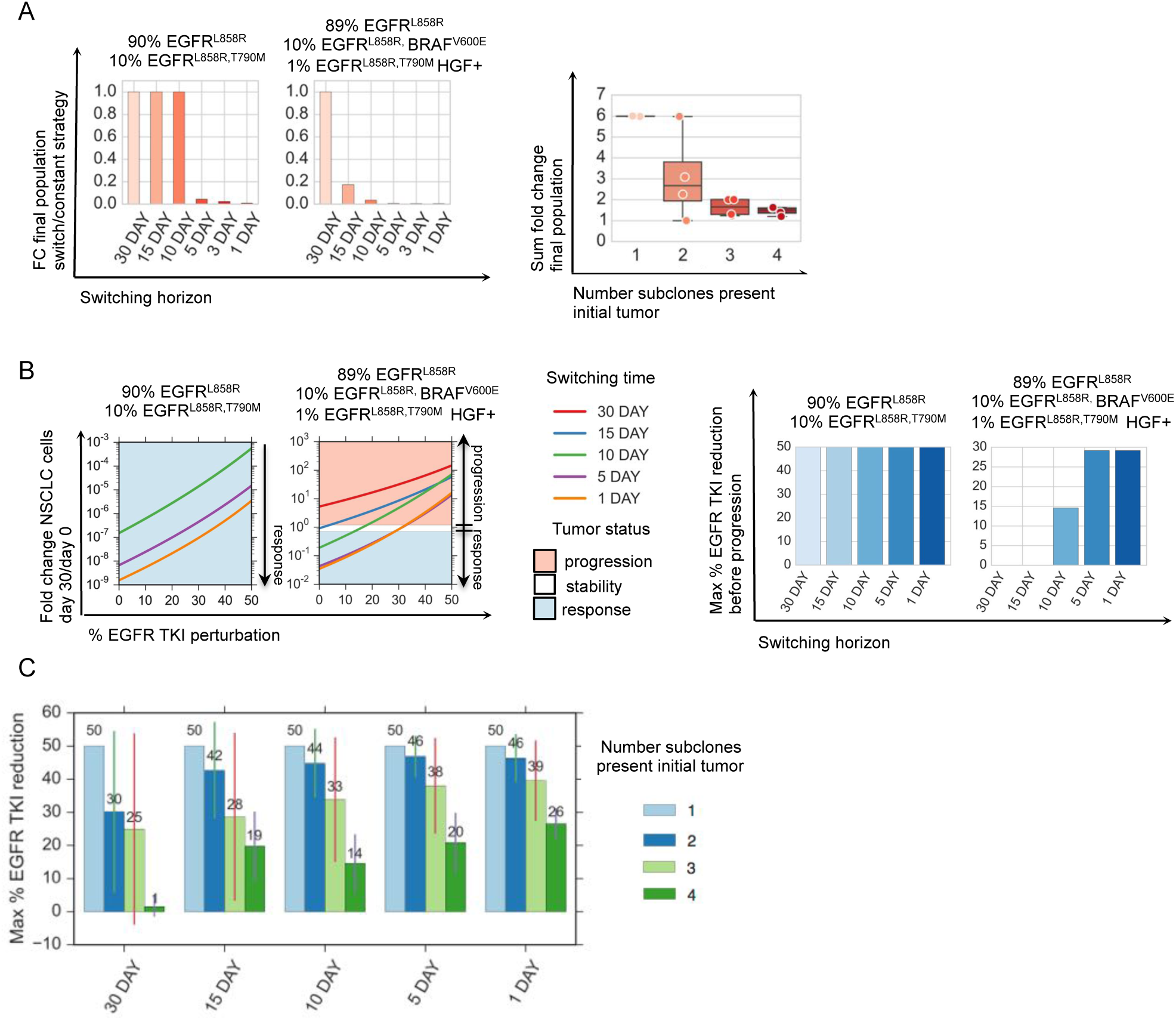
Exploring the robustness of treatment strategies through model simulation. (A) Switching strategies are more beneficial to tumor cell populations with more initial heterogeneity. (Left) Fold change in final lung adenocarcinoma tumor cell populations at day 30 versus day 0 over the course of the optimal 30, 15, 10, 5, 3, and 1 day treatment strategies solved by algorithm 1 (SI, Section 2.2) and normalized by fold change in final tumor cell population for the constant 30 day treatment strategy for an initial tumor cell population comprised of (90% EGFR^L858R^, 10% H1975 EGFR^L858R,T790M^) and another comprised of (89% EGFR^L858R^, 10% BRAF^V600E^,1% EGFR^L858R,T790M^) subclones. (Right) Sum of fold change for the final lung adenocarcinoma populations (SI, Equation S5) for select initial tumor cell distributions (SI, Table 1) and their corresponding optimal 30, 15, 10, 5, 3, and 1 day treatment strategies, categorized by the number of subclones in the initial tumor cell population. Smaller fold change sums indicate that more switching is beneficial to reduce final populations, whereas larger fold changes indicate that more switching does not necessarily help in reducing the final tumor populations. (B) EGFR TKI dose perturbations. (Left) Fold change in number of lung adenocarcinoma cells between day 30 and day 0, as a function of percent EGFR TKI dose reduction for the optimal 30, 15, 10, 5 and 1 day strategies solved by algorithm 1 (SI, Section 2.2) for tumor cell populations indicated above. The shaded areas indicate the regions of the perturbation space where the treatment strategy reduces the initial tumor cell population by more than 30% (response, light blue), increases the size of the original tumor population size by more than 20% (progression, red), or maintains the original tumor population size between the two (stability, white). (Right) Bar graphs indicate the maximum reduction in EGFR TKI dose supported by the optimal strategy such that there is still reduction in tumor size at day 30 with respect to day 0 for the V600E and the pretreatment MET tumor. (C) The average maximum percent EGFR TKI dose reduction supported before progression for lung adenocarcinoma tumors with different number of initial tumor cell subpopulations and for predicted optimal 30, 15, 10, 5, and 1 day switching strategies.

### 3.5 Robustness Analysis of Switching Strategies

Motivated by studies indicating that tissue to plasma ratios for certain drugs such as erlotinib can be low [31], we sought to computationally explore how dose reductions of TKI combinations could affect the evolution of tumor cell populations. This is a particularly relevant clinical issue, as many drugs when used in combination often require a reduction in the recommended monotherapy dose due to toxicity of the dual drug therapy in patients. To examine this question, we simulated the optimal switching strategies corresponding to 30, 15, 10, 5 and 1 day switching horizons subject to EGFR TKI dose reductions for a set of initial tumor cell populations and studied the effects on the final and average tumor populations over the course of the treatment (SI, Mathematical Methods).

For a tumor with a smaller number of initial subclones, such as one comprised of 90% EGFR^L858R^ and 10% EGFR^L858R,T790M^, all switching strategies induced a response for EGFR TKI dose reductions of up to 50% (Fig. 6A). In contrast, with the more complex HGF treated tumor cell population comprised of 89% EGFR^L858R^, 10% EGFR^L858R^BRAF^V600E^, 1% EGFR^L858R,T790M^, only combination strategies with switching horizons of 10 day or shorter induced a response (Fig. 6B). Notably, we observed that the shorter the switching horizon, the higher dose reduction that could be supported while still maintaining a progression free response (Fig. 6B and Fig. S8B). We observed this phenomenon more generally when we simulated different tumor cell initial distributions (Fig. 6C). Thus, we find that the greater number of subclones present in the initial tumor, the greater the benefit there is in increasing switching frequency in terms of the achieving robustness to perturbations in EGFR TKI drug concentration.

### 3.6 Switching Strategies Control or Delay Progression in Vitro

Motivated by the results of our treatment strategy algorithm, we tested drug scheduling strategies on select tumor subpopulations in an *in vitro* model of EGFR mutant lung adenocarcinoma. Specifically, we synthesized the optimal treatment strategy for a heterogeneous HGF treated tumor cell population consisting of 89% EGFR^L858R^, 10% EGFR^L858R^BRAF^V600E^, 1% EGFR^L858R,T790M^, and imposed a constraint that at most one switch could occur, as a starting point to simulate what might be most clinically feasible. The resulting optimal treatment strategy predicted by our modeling, consisting of the erlotinib/crizotinib (days 0-5) followed by the afatinib/trametinib (days 5-30) combination, was shown to elicit the best response *in vitro*, validating our predictive model (Fig. 7B).

**Figure 7:**
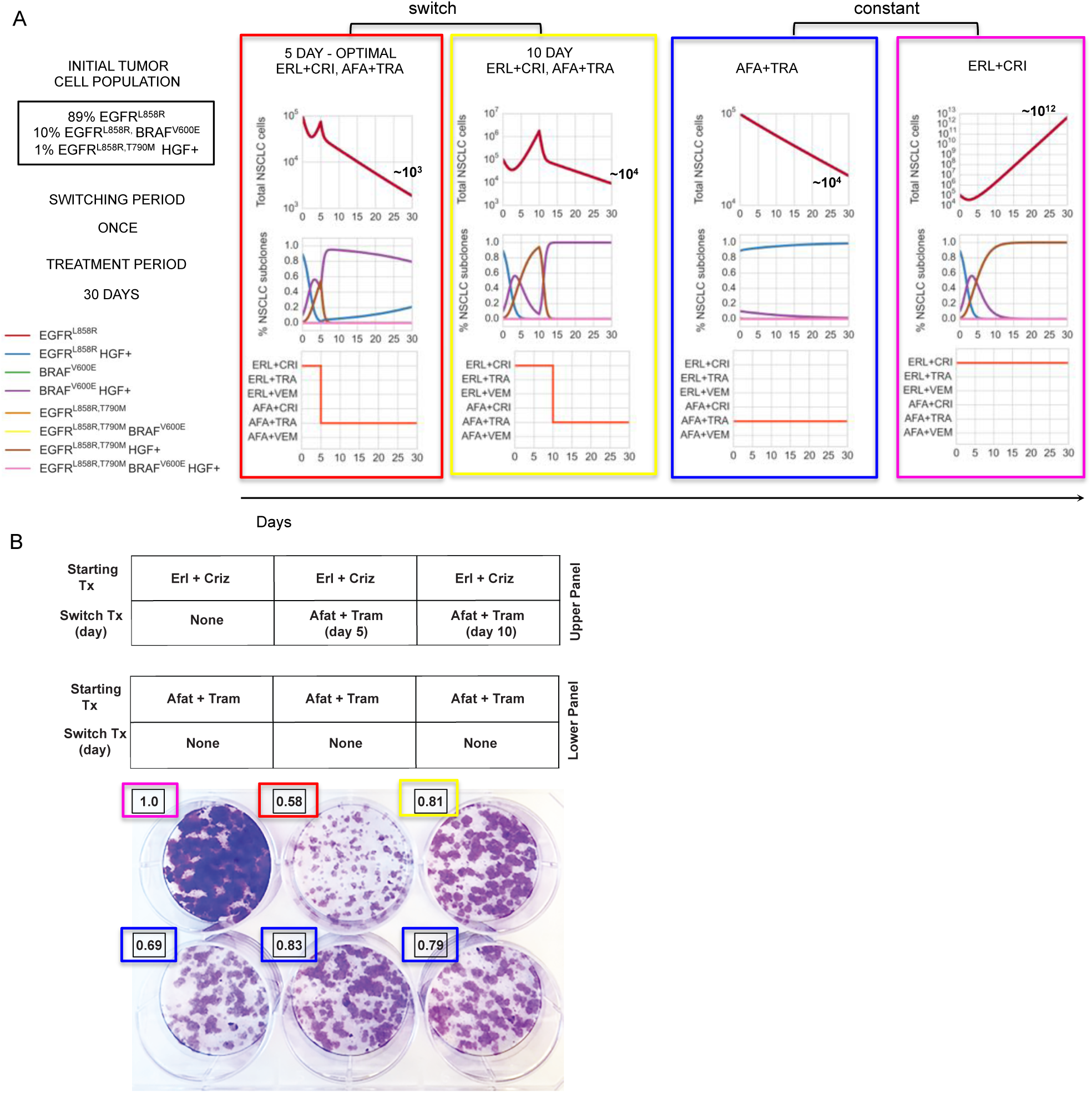
Engineering optimal treatment strategies for concurrent, clonal genetic alterations in EGFR-mutant lung adenocarcinoma and predicting their therapeutic impact. (A) Simulations of the optimal treatment strategy predicted by algorithm 1 (SI, Section 2.2) consisting of 1.5 *µ*M erlotinib+0.5 *µ*M crizotinib for days (0-5) followed by 0.5 *µ*M afatinib+0.5 *µ*M trametinib for days (5-30); the same strategy but with the switch occurring at day 10 and, constant strategies of 0.5 *µ*M afatinib+0.5 *µ*M trametinib or 1.5 *µ*M erlotinib+0.5 *µ*M crizotinib for 30 days, for an initial tumor cell population of 89% EGFR^L858R^, 10% EGFR^L858R^BRAF^V600E^, 1% EGFR^L858R,T790M^, HGF treated. (B) Evolution experiment shows that the predicted strategy for an initial tumor cell population of 89% EGFR^L858R^, 10% EGFR^L858R^BRAF^V600E^, 1% EGFR^L858R,T790M^, treated with 50 ng/ml HGF, is optimal. Overlaid numbers indicate the relative cell density of each well at day 30 compared to the erlotinib+crizotinib well (magenta). Computational simulations in (A) show that the predicted optimal strategy has the greatest reduction in tumor cells in vitro (B, red) compared to the same strategy with a 10 day switch (yellow). A simulation of the model predicts that a constant treatment of afatinib+trametinib produces little change in number of tumor cells (B, blue) and that a constant treatment of erlotinib+crizotinib predicts the exponential outgrowth of the initial EGFR^L858R,T790M^ MET amplified subpopulation, experimentally validated in (B, magenta).

To show how a delay in the switching time might affect response to therapy, we tested equivalent initial tumor cell populations but changed the treatment strategy to start the afatinib/trametinib combination at day 10 instead of at day 5. This resulted in worse overall response than the 5 day switching regimen (Fig. 7B). The corresponding model simulation highlights that although the erlotinib/crizotinib combination effectively targeted the HGF treated EGFR^L858R^ mutation during the first 10 days, it allowed the HGF treated EGFR^L858R, T790M^ subclone to dominate for a longer period of time, thereby impeding overall response.

## 4 Discussion

One of the fundamental challenges in the principled design of combination therapies is the pre-existence and temporal expansion of intratumor genetic heterogeneity that can often lead to rapid resistance with first-line targeted therapies. To address this problem, we sought to develop a new modeling framework to systematically design principled tumor monitoring and therapeutic strategies. We applied a receding horizon optimal control approach to an evolutionary dynamics and drug response model of lung adenocarcinoma that was identified from experimental and clinical data. Based on the clinical and experimental data, our computational method generated optimal drug scheduling strategies for a comprehensive set of initial tumor cell subpopulation distributions.

Our initial insight was that constant drug combination strategies that guarantee progression free response for tumor cell populations with considerable heterogeneity and/or MET activation, required EGFR TKI concentrations that were considerably higher than are typically clinically feasible. At clinically relevant doses, these constant combination strategies were not effective against all tumor cell subpopulations and inevitably, those subpopulations with even slight evolutionary advantages could undergo clonal expansion and cause resistance. To overcome this issue, we used our algorithm to generate optimal drug scheduling strategies that could preempt the outgrowth of these subpopulations over fixed switching periods, and showed that these strategies outperformed constant combination strategies for most tumor cell subpopulation distributions. Notably, our computational analysis showed there was more benefit in applying switching strategies in the context of increasing pre-existing genetic heterogeneity and these switching strategies provided more robustness guarantees in the presence of perturbations in drug concentrations that can occur in patients. We demonstrated successful *in vitro* validation of our optimal control approach for selected tumor subpopulation distributions. In particular, for an *in vitro* analog of our clinical case, a non-intuitive combination therapy switching strategy offered better tumor control than constant treatment strategies.

We found that the most effective drug scheduling strategies were ones that addressed existing subpopulations as they emerged during the course of the treatment, even during a bulk tumor response. In contrast, current standard of care clinical practice is generally to delay switching to second-line therapy until after there is clear evidence of radiographic or clinical progression. Our approach suggests a paradigm shift that would require regular monitoring of an individual patient’s tumor mutational status, for instance by mutational analysis of plasma cell-free circulating tumor DNA, so-called “liquid biopsies” [32, 33, 34, 35, 36]. Our modeling strategy could potentially synthesize this genetic information to yield both the design and prioritization of specific drug regimens and the optimal time for clinical deployment, informed by the molecular findings in a particular patient. Such treatments may need to be applied (non-intuitively) during the initial tumor response, instead of later during therapy or after drug resistance is readily apparent by standard clinical measures in some cases. We envision that our approach could help contribute to the shift from a reactive to pro-active, dynamic management paradigm in solid tumor patients in the molecular era. Drug scheduling strategies synthesized by the algorithm for the initial tumor cell population could be adapted to account for genetic alterations that are detected by the analysis of serial liquid (or tumor) biopsies, leading to a dynamic learning model through iterative refinements; as such, the model could suggest more effective strategies with time. Additional considerations such as pharmacokinetics, the tumor microenvironment and metastatic processes [37, 38] could extend this model to add more clinical relevance. Finally, our approach could guide the optimal timing of serial clinical specimen sampling (plasma, tumor) and radiographic analysis to streamline clinical management. Overall, the combination of techniques stemming from mathematical optimization and control theory combined with more clinically applicable tumor dynamics models is a promising approach to aid the rational design, clinical testing, and clinical adoption of dynamic molecular monitoring and drug scheduling strategies to better control complex solid cancers such as lung cancer in real-time and improve clinical outcomes.

### 4.1 Materials and Methods

#### 4.1.1 Computational Methods

The details of mathematical models and experimental methods may be found in *SI Mathematical Methods*. The mathematical model of lung adenocarcinoma growth mutation and selection by small molecule inhibitors was formulated as system of ordinary differential equations (ODEs). The treatment strategy algorithm was formulated as a receding horizon optimal control problem with the objective of minimizing lung adenocarcinoma populations at every horizon and implemented using python version 3.4.3, scipy version 1.11.0.

#### 4.1.2 Experimental Methods

##### Patient sample preparation and sequence capture

Formalin fixed paraffin embedded (FFPE) NSCLC fine needle aspirate biopsy specimens and a normal blood sample were obtained from the patient under institutional informed consent both prior to erlotinib treatment and upon erlotinib resistance. Lung tumor biopsy specimens contained > 75% tumor cells upon histopathological analysis by a board-certified pathologist. Barcoded sequence libraries were generated using genomic DNA from FFPE tumor material and matched normal blood using the NuGEN Ovation ultralow library systems and according to manufacturer’s instructions (NuGEN, San Carlos, CA). These libraries were among an equimolar pool of 16 barcoded libraries generated and subjected to solution-phase hybrid capture with biotinylated oligonucleotides targeting the coding exons of 389 cancer-associated genes using Nimblegen SeqV.D.J.Cap EZ (Roche NimbleGen, Inc, Madison, WI). Each hybrid capture pool was sequenced in a single lane of Illumina HiSeq2000 instrumentation producing 100bp paired-end reads (UCSF Next Generation Sequencing Service). Sequencing data was demultiplexed to match all high-quality barcoded reads with the corresponding samples.

##### Sequencing Analysis

Paired-end sequence reads from normal blood, pre-treatment tumor, and erlotinib-resistant tumor samples were aligned against build hg19 of the reference genome with BWA [39]. Duplicate reads were marked, alignment and hybridization metrics calculated, multiple sequence realignment around candidate indels performed, and base quality scores recalibrated across all samples with the Picard suite (http://picard.sourceforge.net/) and the Genome Analysis Toolkit (GATK) [40]. Somatic point mutations were detected in the treatment-naïve and resistant tumors using MuTect [41], while small insertions and deletions (indels) were identified with GATK. Given the depth of sequencing achieved and the presence of low-frequency oncogenic mutations in the normal sample likely due to circulating tumor DNA, mutations were excluded as germline if they exceeded a frequency of 10% in the normal sample. Non-synonymous mutations were annotated for their sequence context, effect, and frequency in lung adenocarcinoma and squamous cell tumors from The Cancer Genome Atlas (TCGA) project and Imielinski et al. [42]. All previously characterized oncogenic alleles in NSCLC or mutations previously linked to erlotinib resistance were also manual inspected in both treatment-naïve and resistant tumors. This analysis revealed a single sequencing read bearing the T790M mutation in the primary tumor (total coverage at this locus: 1300x). This was insufficient evidence from sequencing data to formally call the mutation, but we cannot exclude the possibility that EGFR^T790M^ exists pre-treatment in a very rare clone (<0.08%) for which our target depth of coverage limited our sensitivity. DNA copy number alterations where inferred from the mean sequence coverage for each target region in each sample corrected for overall library size. Amplifications and deletions were determined from ratios of coverage levels between the pre- and post-treatment tumors and the matched normal blood sample. Due to the elevated signal to noise from targeted capture and sequencing of FFPE material from lower input amounts, overt genomic amplifications and deletions were required to affect multiple target regions (exons) of a given gene before being called as detected. The EGFR^T790M^ and BRAF^V600E^ variants were confirmed by a standard Clinical Laboratory Improvement Amendments (CLIA)-approved PCR-based shifted termination assay (data not shown). The changes in EGFR and MET copy number were validated using established fluorescence in situ hybridization clinical assays.

##### Cell Lines and Reagents

Human lung cancer cell lines were acquired as previously described [43, 44]. Cells were grown in RPMI 1640 supplemented with 10% (high serum) or 0.5% (low serum) fetal bovine serum (FBS), penicillin G (100U/ml) and streptomycin SO4 (100U/ml). Erlotinib, afatinib, vemurafenib, crizotinib, and trametinib were purchased from Selleck Chemicals (Houston, TX). Drugs were resuspended in DMSO at a concentration of 10mM and stored at −20 °C. Erlotinib and afatinib were used at working concentrations ranging from 0.010-1.5 *µ*M. Vemurafenib was used at a working concentration of 5.0 *µ*M, and trametinib and crizotinib were used at a 0.5 *µ*M. HGF was purchased from Peprotech (Rocky Hill, NJ) and resuspended at 50 g/ml in sterile PBS + 0.1% BSA. Cells were treated with HGF at 50 ng/ml.

##### Generation of stable cell lines

293-GPG viral packaging cells were transfected with pBABE (empty vector), pBABE-mCherry-BRAF-WT and pBABE-mCherry-BRAFV600E constructs (kindly provided by Dr. Eric Collision, UCSF, San Francisco, CA) using Lipofectamine-2000 (Life Technologies, Pleasanton, CA) per manufacturer’s instructions. Virus containing media was harvested three days post transfection and used to infect 11-18 and H1975 lung cancer cell lines. Cells were incubated with virus containing media supplemented with 6 *µ*g/ml of polybrene for 24 hours. Media was changed to standard cell growth media (RPMI-1640 + 10% fetal bovine serum and 100 U/ml penicillin G and 100 U/ml streptomycin SO4) and cells were expanded for 48 hours, at which point puromycin (2 *µ*g/ml) was added to the media and cells were allowed to grow for an additional 4 days. Cells that survived puromycin selection (stable cell lines) were used in all subsequent experiments.

##### Cell Viability and Growth Assays

Assays were performed as previously described [43, 44]. Briefly, cells were seeded overnight at a density of 5,000 cells per well in 96-well plates in RPMI containing 10% FBS and treated with indicated reagents for 72 hours. Viable cell numbers were determined using the CellTiterGLO assay according to manufacturer’s instructions (Promega). Each assay consisted of six replicate wells and was repeated at least twice in independent experiments. Cell viability is presented as the mean (*±* s.e.m.) erlotinib or afatinib inhibitory concentration 50 (IC50). Statistical significance between treatment groups was determined by the Bonferroni’s multiple comparisons ANOVA statistical test.

##### Immunoblot analysis

Cells were harvested 24h after initiation of treatment with reagents. Cells were scraped and lysed in lysis buffer (50 mM Tris*·*HCl pH 8.0, 150 mM sodium chloride, 0.1% SDS, 0.5% sodium deoxycholate, 1% Triton X 100, 5 mM EDTA containing protease and phosphatase inhibitors (Roche Diagnostics. Indianapolis, IN). After quantitation by Pierce BCA assays (Thermo Scientific, Rockford, IL), 25 *µ*g of each sample was separated by gel electrophoresis on 4-15% Criterion TGX precast gels (BioRad, Hercules, CA) and transferred to nitrocellulose membrane. For immunoblots, the following antibodies were used: anti-total EGFR (1:1000 dilution, Bethyl Laboratories, Inc., Montgomery TX), anti-pEGFR, anti-total Met, anti-pMet, anti-total Mek, anti-pMek, anti-total Akt, anti-pAkt, anti-total Erk, anti-pErk (1:1000, Cell Signaling Technology Inc., Danvers, MA), BRAF^V600E^ Monoclonal Antibody (Clone VE1, 1:1000, Spring Bio-science, Pleasonton, CA), BRAF WT (1:1000, Santa Cruz Biotech, Santa Cruz, CA) and anti-actin (1:5000 dilution, Sigma-Aldrich, Saint Loius, MO), HRP-conjugated anti-rabbit Ig (used at a 1:3000 dilution, Cell Signaling), and HRP-conjugated anti-mouse IgG (used at a 1:3000 dilution, Cell Signaling). Specific proteins were detected by using either ECL Prime (Amersham Biosciences, Sunnyvale, CA) or the Odyssey Li-Cor (Lincoln, NE) with the infrared dye (IR Dye 800, IR Dye 680)-conjugated secondary antibodies (1:20,000, Li-Cor).

## 5 Acknowledgements

The authors acknowledge funding support from NIH, the Pew-Stewart Charitable Trust, the Kinship-Searle Foundation, and the Van Auken Foundation (to TGB). C.M.B was supported by grants from The Lung Cancer Research Foundation (P0060520) and AACR (14-40-18-BLAK). J.C.D. provided funding for V.D.J. and N.M..

## 6 Author contributions

T.G.B.,V.D.J. and C.M.B. conceived and designed the study. V.D.J. conceived and developed the math model and algorithm. V.D.J. and N.M. implemented the algorithm. C.M.B and V.D.J., designed experiments, performed experiments and analyzed data. V.O. and L.L. performed experiments. L.L., and B.C.B. performed sequencing. S.A., B.C.B., and B.S.T. analyzed sequencing data. M.A.G. provided tumors for analysis. V.D.J., C.M.B. and T.G.B. wrote the manuscript, with input from all authors. We thank Nikoletta Sidiropoulos for pathology assessment and independent confirmation of the BRAF V600E mutation. We thank Tyrrell Nelson for assistance in sequencing library preparation.

## Supplementary Information

### 1 Mathematical Methods

#### 1.1 Evolutionary Dynamics Model of NSCLC

The quasispecies model [1] was originally developed to describe the dynamics of populations of self replicating macromolecules undergoing mutation and selection. We choose this model for its relative simplicity and its ability to capture the salient features of the evolutionary dynamics of a simplified generic disease model. The following adaptation incorporates the effects of small molecule inhibitors and describes the growth, mutation and evolution of non small cell lung adenocarcinoma populations:

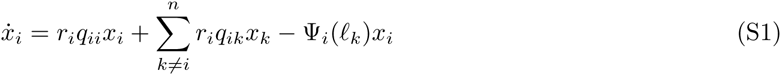

where *x*_*i*_ *ε* ℝ_+_ is the concentration of a NSCLC subpopulation *i*, *ℓ*_*k*_ *ε* ℝ_+_ is a small molecule inhibitor concentration (assumed to remain at constant concentrations throughout), *r*_*i*_ is the growth rate for each cell *x*_*i*_, and *q*_*ik*_ is the probability that cell *k* mutates to cell *i* (note that *q*_*ii*_ is the probability of no mutation occurring). Finally, the function Ψ_*i*_(*ℓ*_*k*_) represents the pharmacodynamics of individual drugs *ℓ*_*k*_ or of individual EGFR TKIs (erlotinib or afatinib) in combination with fixed concentrations other small molecule inhibitors used in this study (0.5 *µ*M crizotinib, 0.5 *µ*M trametinib or 5 *µ*M vemurafenib) with respect to the *i*-th NSCLC cell type, namely:

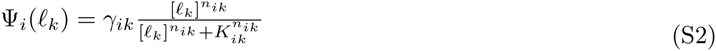

where *ℓ*_*k*_ *ε* ℝ_+_ is the drug concentration, *γ*_*ik*_ *ε* ℝ_+_ is the saturation coefficient, *K*_*ik*_ *ε* ℝ_+_ is the dissociation constant, *n*_*k*_ *ε* ℝ_+_ is the Hill coefficient. When *ℓ*_*k*_ = 0, ∀*k ε* {1, …, *m*}, the dynamics are unstable.

### 2 A control theoretic algorithm for designing treatment strategies

To design treatment strategies that best minimize tumor size and control its evolution over time, we combine both a greedy algorithm and receding horizon control approach. We introduce some notation, cost function definitions and specify our algorithm.

#### 2.1 Cost functions

To measure the effectiveness of a given treatment strategy over time, we define the *average cost* function. For a given treatment strategy *ℓ*_*k*_ applied to Equation (S1), we rewrite the dynamics of the entire system

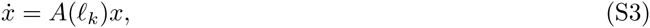

where *A ε* ℝ^*n*×*n*^ is a matrix that represents the growth, mutation and drug dynamics for treatment strategy *ℓ*_*k*_, for *n* cell subpopulations.

The *average cost C*_*r*_ for a time horizon *N*, allowable switching period *τ* and time intervals of the form [*kτ*, (*k* + 1)*τ*] for *k* = {0, .., *N*/*τ* − 1} is given by

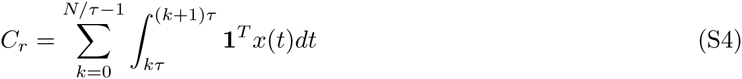

where 1*^T^* is the *n* × 1-dimensional vector of ones and *x*(*t*) is the solution to Equation (S3).

Equation (S4) simplifies to

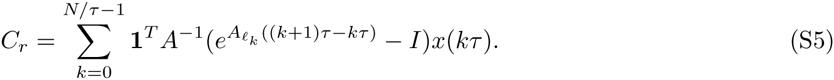

The *final cost C*_*f*_ for an inital tumor population *x*(0) and a sequence of drugs 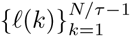 that define a switching therapy over a time horizon *N* is defined as

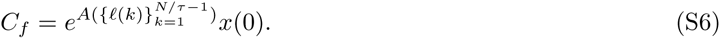

#### 2.2 Algorithm

Our algorithm is defined as follows. Given an initial tumor population, denoted by *x*_0_, a time horizon *N* and an allowable switching period *τ*, we perform the following computations to determine a candidate treatment strategy:

##### Algorithm 1 Treatment strategy synthesis

1. **Initialization:** Set *k* = 0 and *x*(0) = *x*_0_.
2. **Greedy approach:** For time interval [*kτ*, (*k* + 1)*τ*], compute 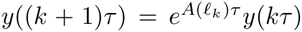 for each possible treatment strategy *ℓ*_*k*_.
3. **Update:** Set 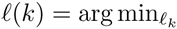 sum(*y*(*k* + 1)*τ*), and set 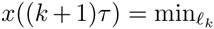 sum(*y*(*k* + 1)*τ*). Increment *k*: if *k* = *N*, proceed to step 4, otherwise return to step 2.
4. **Output:** A sequence of drugs 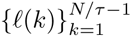 that define a switching therapy.

The resulting switching therapy {*ℓ*(*k*)} is then applied until the next biopsy can be taken, giving a new tumor cell population measurement, at which point the algorithm is repeated. In particular, it is important that the horizon *N* be chosen to be longer than expected periods between biopsies.

### 3 Model Implementation and Simulations

#### 3.1 Derivation of dynamical system parameters

**Growth and Mutation Rates**. We model the growth of NSCLC cell population *x*_*i*_ by the following ordinary differential equation (ODE):

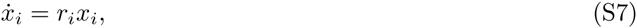

where *r*_*i*_ is the growth rate per day, and 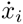 denotes the derivative with respect to time of the tumor cell population *x*_*i*_. Note that we assume that no mutations occur over the time-frame considered, allowing us to set *q*_*ii*_ = 1 and *q*_*ij*_ = 0 in the dynamic model (S1), resulting in (S7).

Given an initial population *x*_*i*_(0), the population *x*_*i*_(*t*) on day *t* can be obtained by solving ODE (S7), and is specified by the following expression

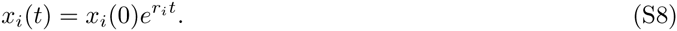

Given a set of *N* experimental data points *e*_*i*_(0), *e*_*i*_(*t*_1_), …, *e*_*i*_(*t*_*N*_), we fit these points to an exponential function of the form (S8), with *x*_*i*_(0) = *e*_*i*_(0) to obtain an experimentally derived value for the growth rate *r*_*i*_ of tumor cell population *x*_*i*_.

We take the DNA mutation rate to be 1*e*^−9^ mutation/base pair/cell division []. We assume that mutations occur unidirectionally from EGFR^L858R^ parental cells to EGFR^L858R,T790M^, EGFR^L858R^, BRAF^V600E^ or EGFR^L858R,T790M^ BRAF^V600E^, HGF-/+.

##### Drug Effect Rates and Hill Functions

We model the change in a tumor cell population *x*_*i*_ under a treatment *j* of concentration *ℓ* with the following ordinary differential equation (ODE):

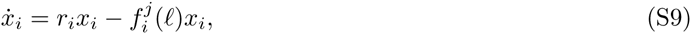

where *r*_*i*_ is the growth rate per day derived in the previous section and 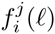 is a function mapping the treatment *j* at concentration *ℓ* to a drug effect rate per day. We again assume that no mutation occurs over the time-frame considered, allowing us to set the mutation rates *q*_*ii*_ = 1 and *q*_*ij*_ = 0 in the model (S1), resulting in (S9).

Similar to the previous section, given an initial population *x*_*i*_(0), the population *x*_*i*_ (*t*) on day *t* can be obtained by solving ODE (S9), and is specified by the following expression.

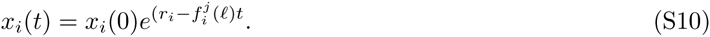

We model the map 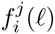 as a modified function of the form

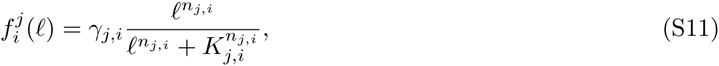

where *γ*_*j,i*_ *n*_*j,i*_ and *K*_*j,i*_ are the saturation parameter, Hill function coefficient and binding reaction dissociation constant for drug *j* applied to cell *x_i_*.

Our goal is to obtain values for these three parameters using experimental data measuring cell viability under varying concentrations *ℓ* of drug *j*. In particular, given experimentally obtained data pairs of the form *ℓ*, *y*_*i,j,ℓ*_(1), where *y*_*i,j,ℓ*_(1) is the ratio of the tumor cell population *x_i_* treated with concentration *ℓ* of drug *j* at day 1 to the tumor cell population *x*_*i*_ treated with no drug at day 1. Letting 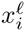 denote the treated tumor population and 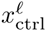 denote the untreated control tumor population, it follows that *y*_*i,j,ℓ*_ can be written as

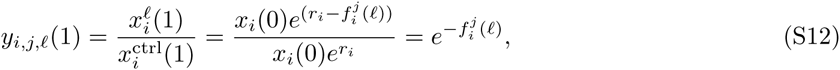

where the first equality follows from the definition of *y*_*i,j,ℓ*_(1), the second from applying equations (S10) and (S8) to 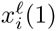 and 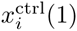 respectively, and the third from canceling like terms. It follows that the experimentally derived values of 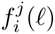 are given by

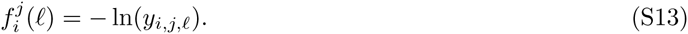

Solving this equation for each experimentally tested concentration *ℓ*, we obtain a set of points 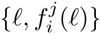 that can be used to derive the parameters *γ*_*j,i*_ *n*_*j,i*_ and *K*_*j,i*_ via curve fitting. In order to avoid overfitting, we set 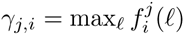, i.e., we force the modified Hill function to saturate at the maximal experimentally observed rate. Although this approach can be conservative in modeling the drug effect rate of high concentrations of drugs, we note that the the maximal dose tested is chosen to be significantly higher than the maximum tolerated doses, and hence we do not expect this saturation to affect the accuracy of our model at clinically relevant doses.

#### 3.2 Evolutionary stability measured by maximum eigenvalues

Figures (S7) and (main text) depict maximum eigenvalue decompositions of HGF- and HGF+ tumors and describe the set of initial NSCLC populations, if present can lead to tumor progression upon initiation of constant (non-switching) combination treatments. For the evolutionary dynamics:

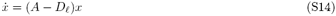

where *x ε* ℝ^*n*^ is a vector of concentrations of *n* NSCLC subpopulations, 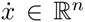 is their rate of change over time, *A ε* ℝ^*n*×*n*^ is a matrix that represents the growth and mutation dynamics and *D*_*ℓ*_ *ε* ℝ^*n*×*n*^ is a diagonal matrix that represents the corresponding drug effect of one constant drug treatment on the rate of change of NSCLC cells. If all eigenvalues are negative then Equation (S14) is said to be *stable*. In the case of NSCLC evolutionary dynamics corresponding to Equation (1), *stability* refers to tumor reduction, and *instability* refers to tumor progression. In section 3.1, we made the assumption that mutation rates are one directional, hence the *A* matrix in Equation (1) is lower triangular and the eigenvalues of *A* − *D*_*ℓ*_ are exactly equal to its diagonal entries. For each NSCLC subpopulation, we take the maximum eigenvalue for each evolutionary branch downstream of the population and define this as *evolutionary stability*. This maximum eigenvalue represents the worst case stability if the particular population is present upon treatment initiation - a positive maximum eigenvalue indicates that the presence of the cell subpopulation in the tumor upon initiation of treatment is likely to cause therapeutic failure. A negative maximum eigenvalue indicates that the presence of the particular subpopulation will not outgrow or evolve in the presence of therapy.

#### 3.3 Robustness analysis

##### Sensitivity to drug perturbations

To analyze the effect of dose reductions on the robustness of constant and switching treatment strategies, we perturbed the drug concentrations and calculated the ratio of final cost and initial cost (Figures (S8)). We rewrite Equation (S1) for one cell *x*_*i*_ and one drug *ℓ*_*j*_ to illustrate how a drug perturbation *δ ε* ℝ^[0,1]^ is modeled:

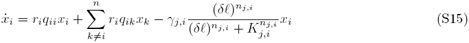

The fold change *F C*_*f*_ in total population from day 0 to day *N* for a sequence of drugs 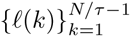 defining a switching strategy over a time horizon *N*, and initial tumor population *x*_0_ = *x*(0) is calculated by

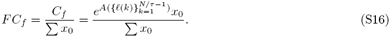

If *F C*_*f*_ < 1, the treatment strategy 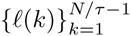 is effective for NSCLC populations for the duration of the time horizon *N*, *F C*_*f*_ > 1 indicates progression.

#### 3.4 Implementation

The evolutionary dynamics model and simulations were implemented using python, scipy and numpy (versions 3.5.1, 0.17.0, 1.9.3) and pandas version 0.17.0 was used for data parsing. Data fitting for experimentally derived cell growth and drug dose response data was performed with Matlab version 8.3.0.532 using the non linear least squares method.

**Figure S1:**
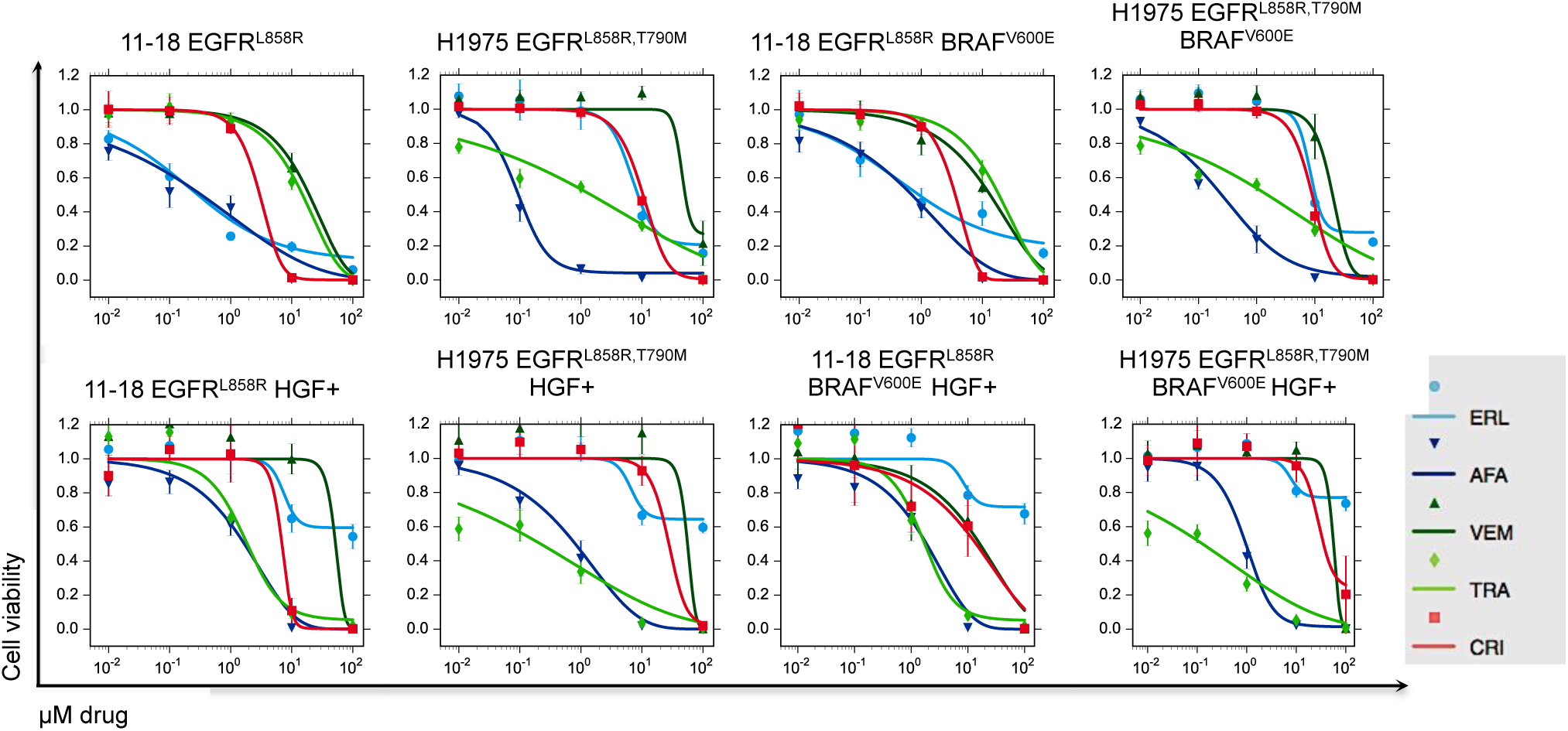
Experimentally derived erlotinib, afatinib, vemurafenib, trametinib and crizotinib dose response curves for 11-18 EGFR^L858R^, 11-18 EGFR^L858R^ BRAF^V600E^, H1975 EGFR^L858R,T790M^ H1975 EGFR^L858R,T790M^ BRAF^V600E^ cell lines, and either 0 or 50 ng/ml human growth factor (HGF) and fit with 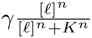 where *γ* is the maximum inhibition, [*ℓ*] is the EGFR TKI concentration, *n* is the Hill coefficient and *K* is the half maximal inhibitory concentration (IC50).

**Figure S2:**
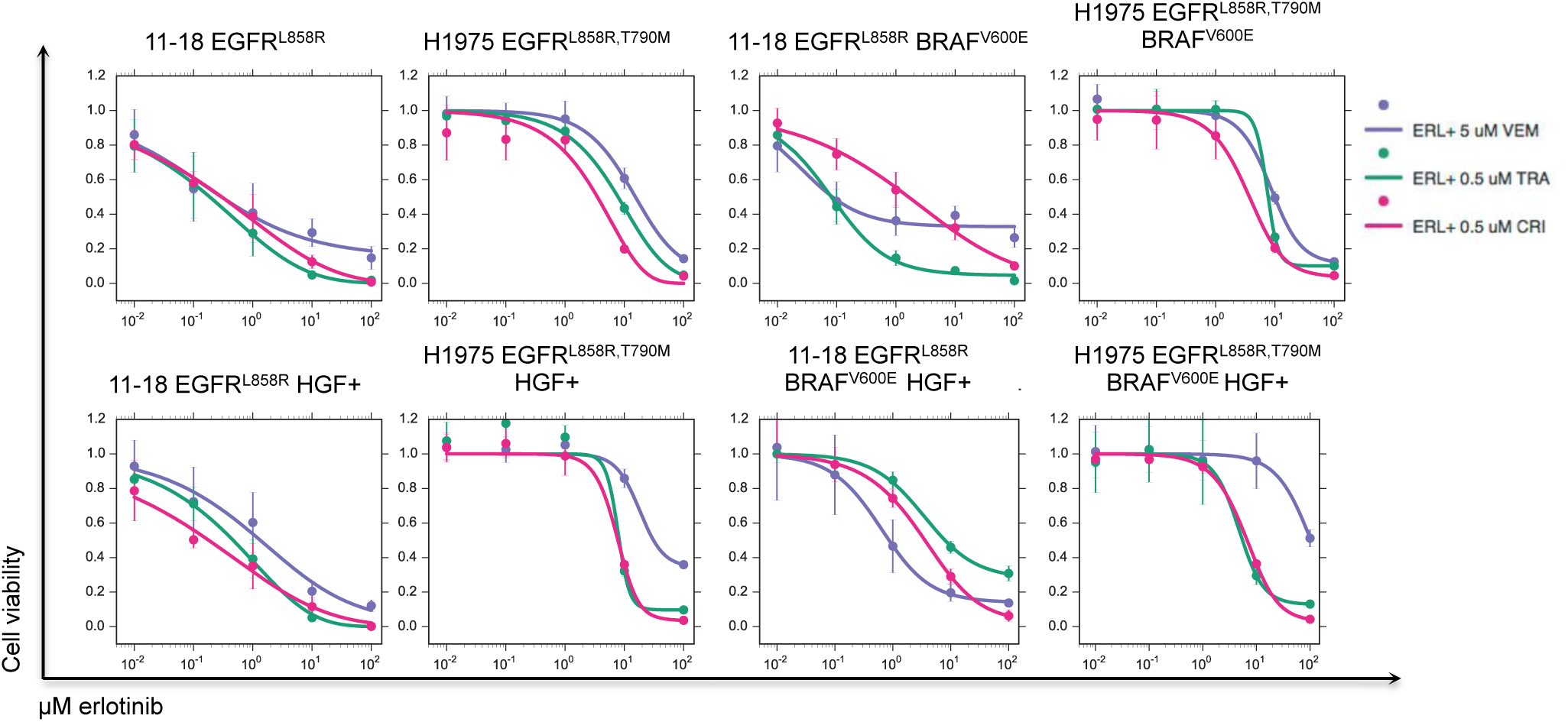
Experimentally derived dose response curves for erlotinib in combination with 5 *µ*M vemurafenib, 0.5 *µ*M trametinib and 0.5 *µ*M crizotinib for 11-18 EGFR^L858R^, 11-18 EGFR^L858R^BRAF^V600E^, H1975 EGFR^L858R,T790M^ H1975 EGFR^L858R,T790M^ BRAF^V600E^ cell lines, and either 0 or 50 ng/ml human growth factor (HGF) and fit with 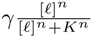 where *γ* is the maximum inhibition, [*ℓ*] is the EGFR TKI concentration, *n* is the Hill coefficient and *K* is the half maximal inhibitory concentration (IC50).

**Figure S3:**
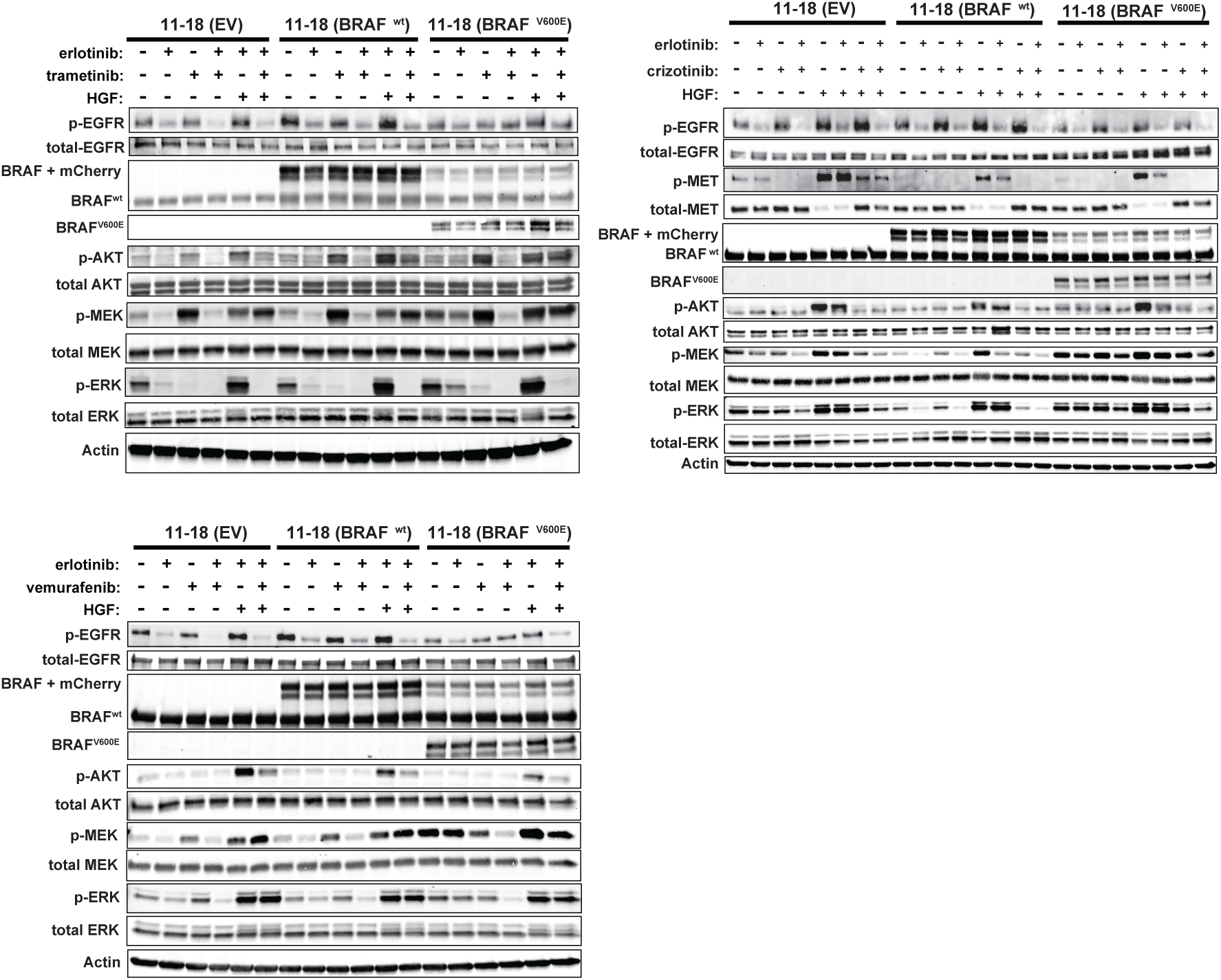
Western blot analysis of cell lysates obtained from 11-18 cell line, treated with drugs and/or HGF as indicated, and probed for the indicated proteins.

**Figure S4:**
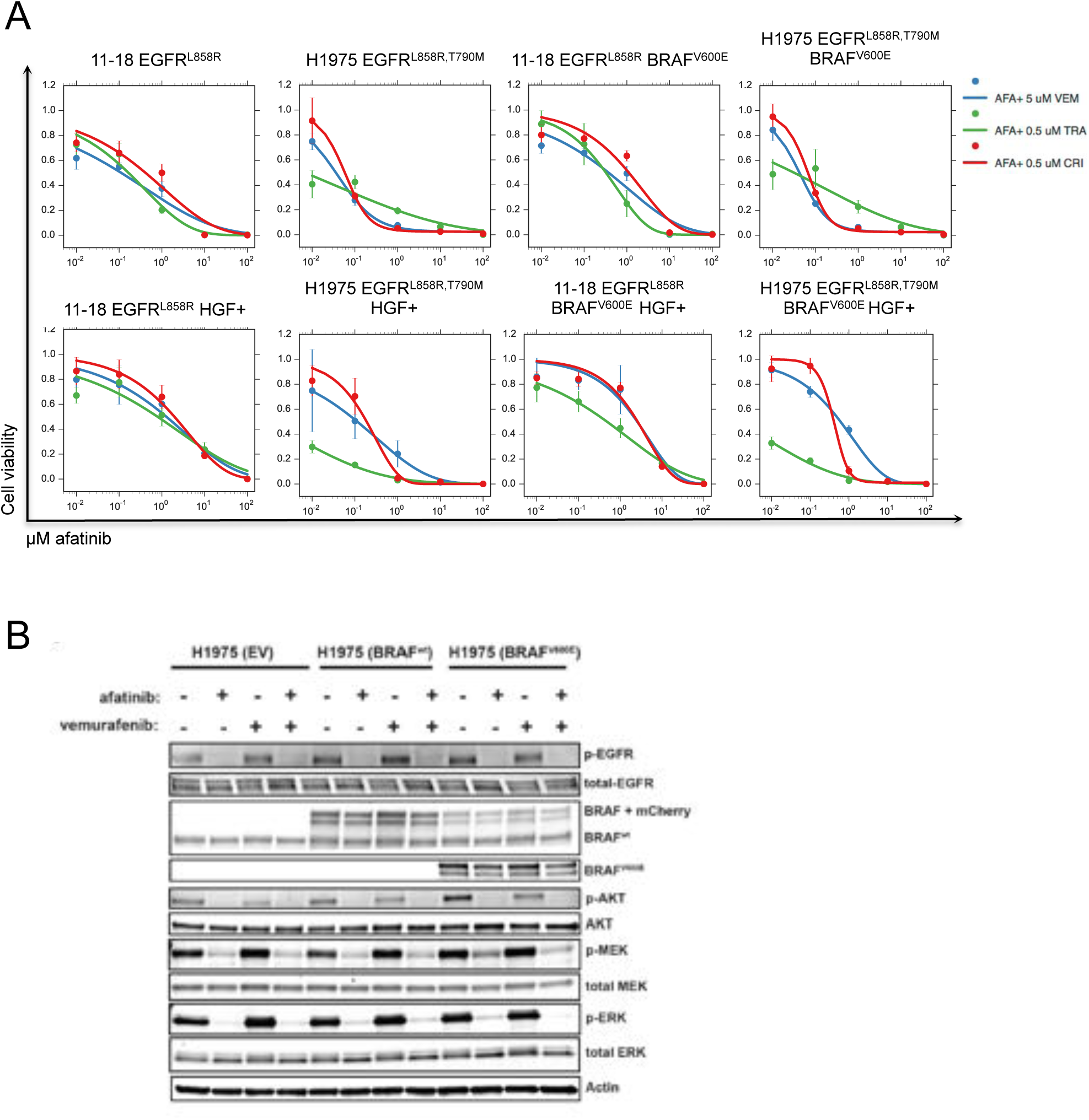
A) Experimentally derived dose response curves for afatinib in combination with 5 *µ*M vemurafenib, 0.5 *µ*M trametinib and 0.5 *µ*M crizotinib for 11-18 EGFR^L858R^, 11-18 EGFR^L858R^ BRAF^V600E^, H1975 EGFR^L858R,T790M^ H1975 EGFR^L858R,T790M^ BRAF^V600E^ cell lines, and either 0 or 50 ng/ml human growth factor (HGF) and fit with 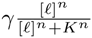 where *γ* is the maximum inhibition, [*ℓ*] is the EGFR TKI concentration, *n* is the Hill coefficient and *K* is the half maximal inhibitory concentration (IC50). (B) Western blot analysis of cell lysates obtained from H1975 cell lines, treated with drugs and/or HGF as indicated, and probed for the indicated proteins.

**Figure S5:**
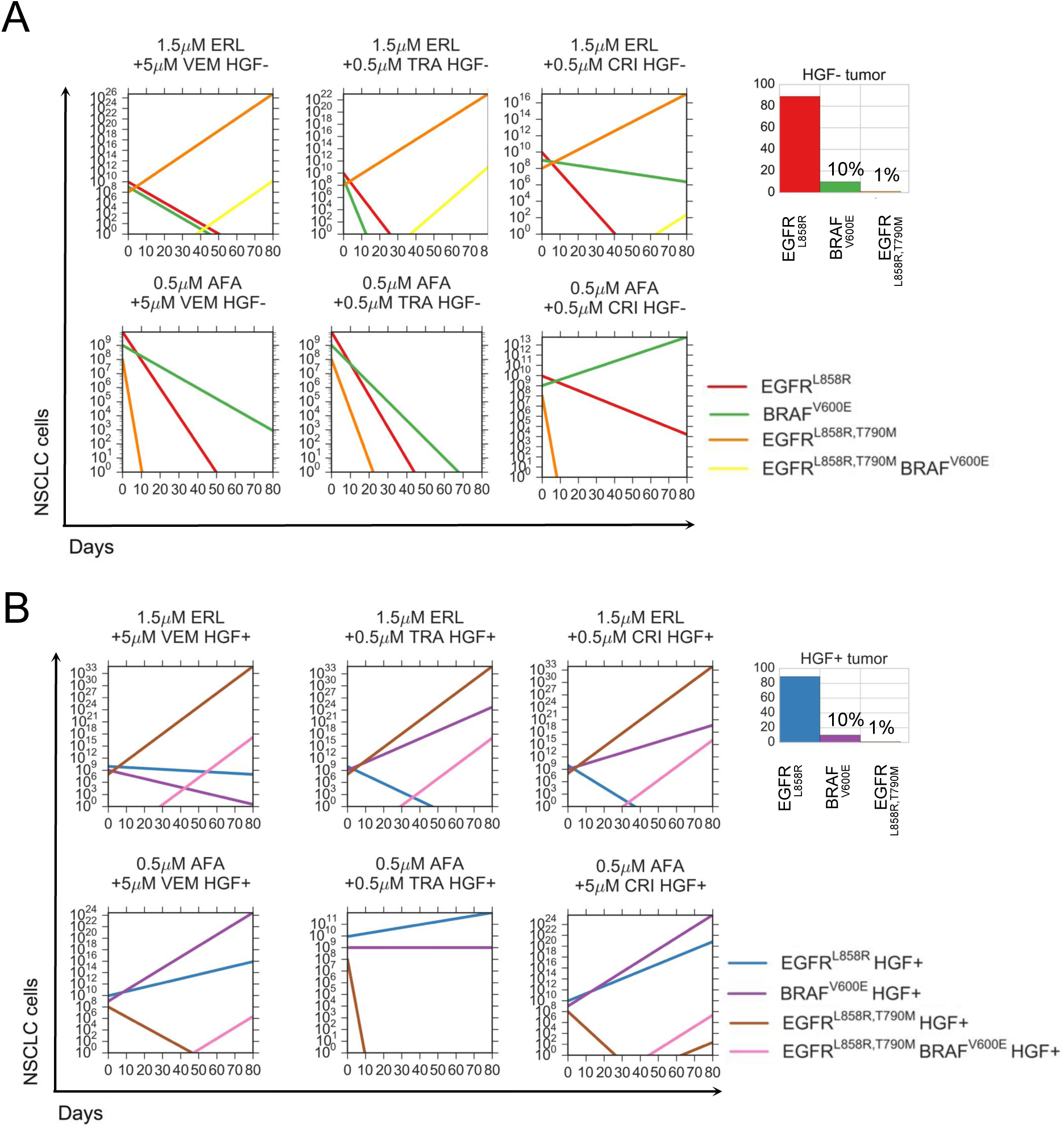
Simulations of the NSCLC model for constant combinations of 0.5 *µ*M afatinib or 1.5 *µ*M erlotinib with either 0.5 *µ*M trametinib, 0.5 *µ*M crizotinib or 5 *µ*M vemurafenib for a tumor comprised of 89% 11-18 EGFR^L858R^, 10% 11-18 EGFR^L858R^, BRAF^V600E^ and 1% H1975 EGFR^L858R T790M^, and treated with HGF (B) or without HGF

**Figure S6:**
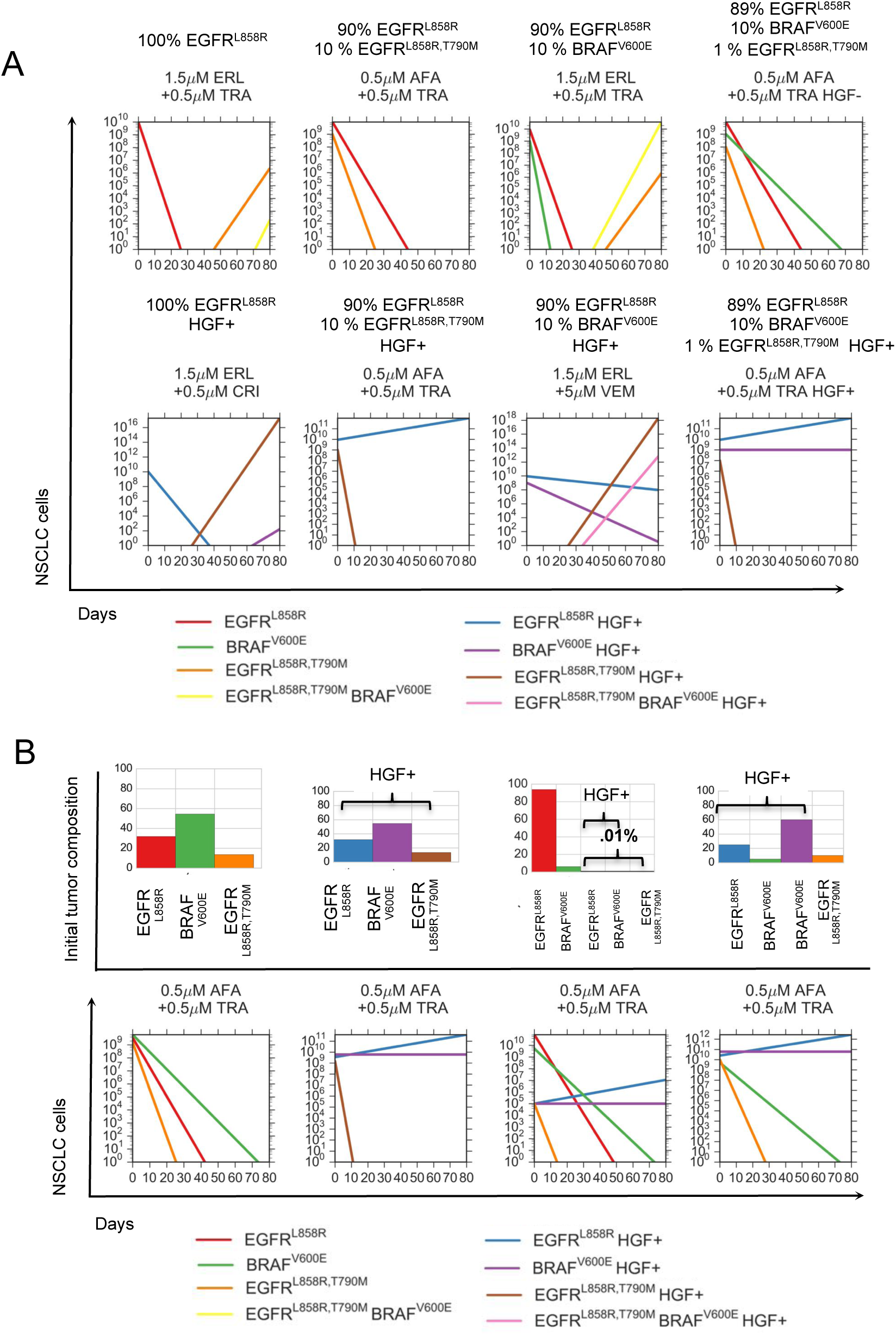
Simulations of the NSCLC model for the optimal 30 day constant combinations found by Algorithm (4) with 0.5 *µ*M afatinib or 1.5 *µ*M erlotinib with either 0.5 *µ*M trametinib, 0.5 *µ*M crizotinib or 5 *µ*M vemurafenib for the relatively low (A) initial tumor heterogeneity or with (B) high initial tumor heterogeneity.

**Figure S7:**
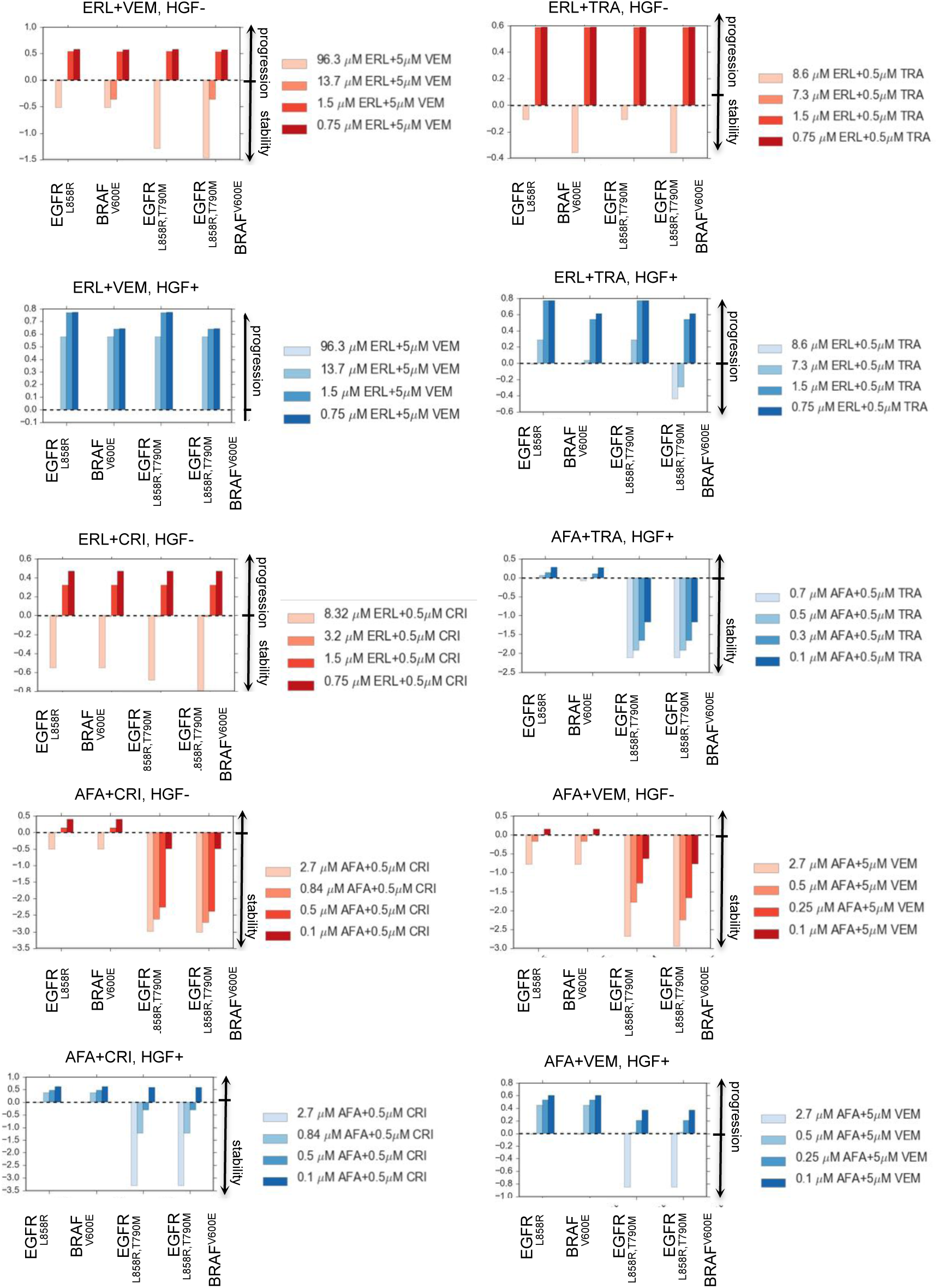
Classification of initial tumor compositions via eigenvalue decompositions describe the initial tumor populations that can destabilize of the evolutionary dynamics in the presence of either erlotinib or afatinib and either 0.5 *µ*M trametinib, 0.5 *µ*M crizotinib or 5 *µ*M vemurafenib.

**Figure S8:**
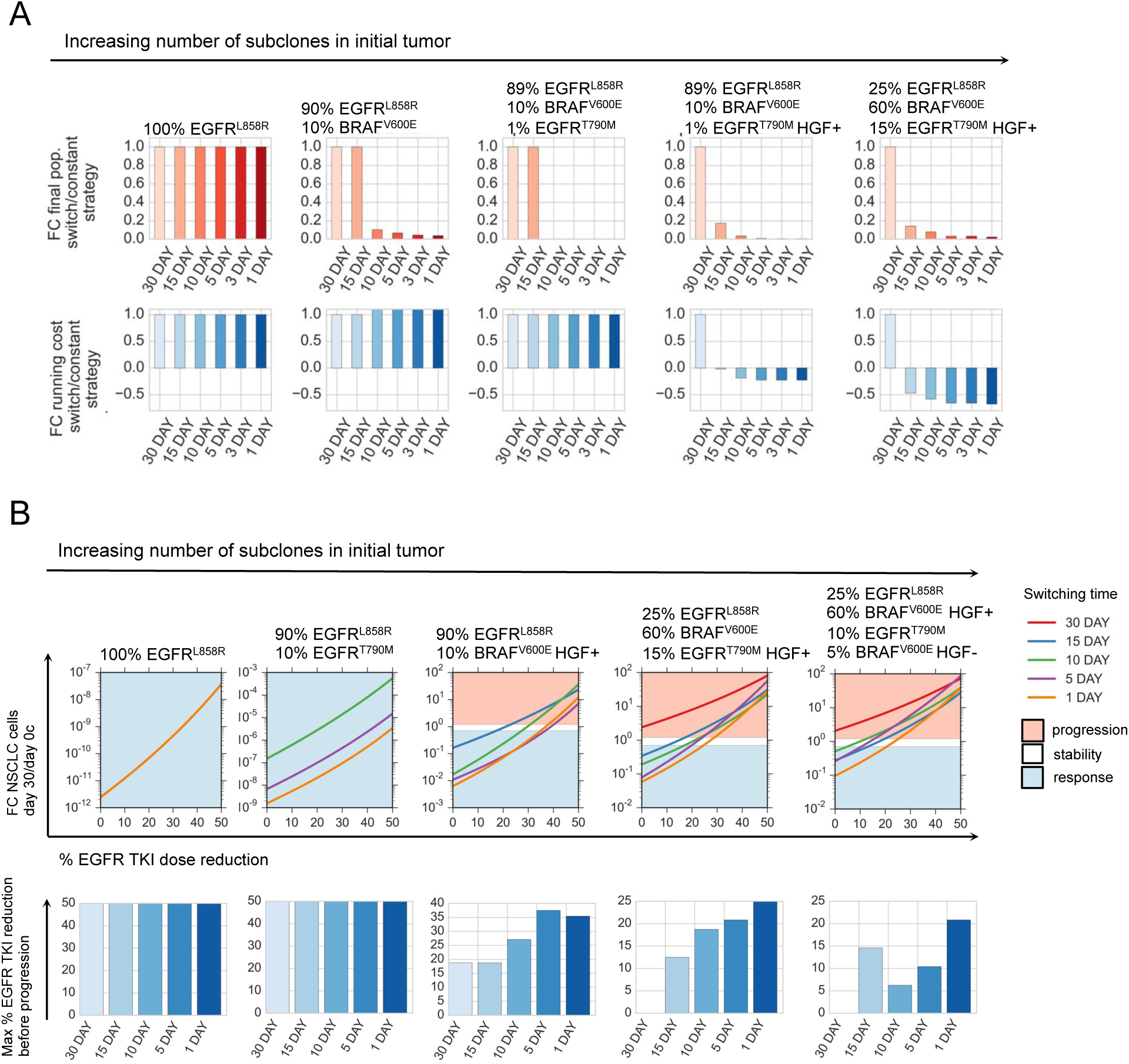
A) Fold change in NSCLC population at day 30 versus day 0, over the course of the optimal 30, 15, 10, 5, 3, and 1 day treatment strategies solved by algorithm 1 (SI), for indicated tumor compositions, normalized by fold change in NSCLC population for the constant 30 day treatment strategy (Red). (Blue) Sum of fold change in the average cost for indicated tumor compositions and corresponding optimal 30, 15, 10, 5, 3, and 1 day treatment strategies. B) (Above) Fold change in number of NSCLC cells between day 0 and day 30, as a function of percent EGFR TKI dose reduction for the optimal 30, 15, 10, 5 and 1 day strategies solved by algorithm 1 (SI) for indicated tumor compositions. Shaded blue areas indicate the region of the perturbation space where the treatment strategy reduces the size of the initial tumor (stable). The shaded red area indicates the region of the perturbation space where the treatment strategy increases the size of the original tumor at day 30 (unstable). (Below) The maximum percent EGFR TKI dose reduction sustainable before the treatment is no longer effective (the tumor progresses).

**Figure S9:**
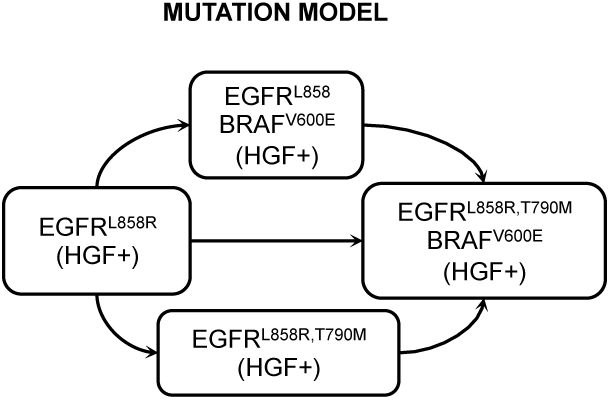
The EGFR-mutant lung adenocarcinoma mutation model used in this study.

**Figure S10:**
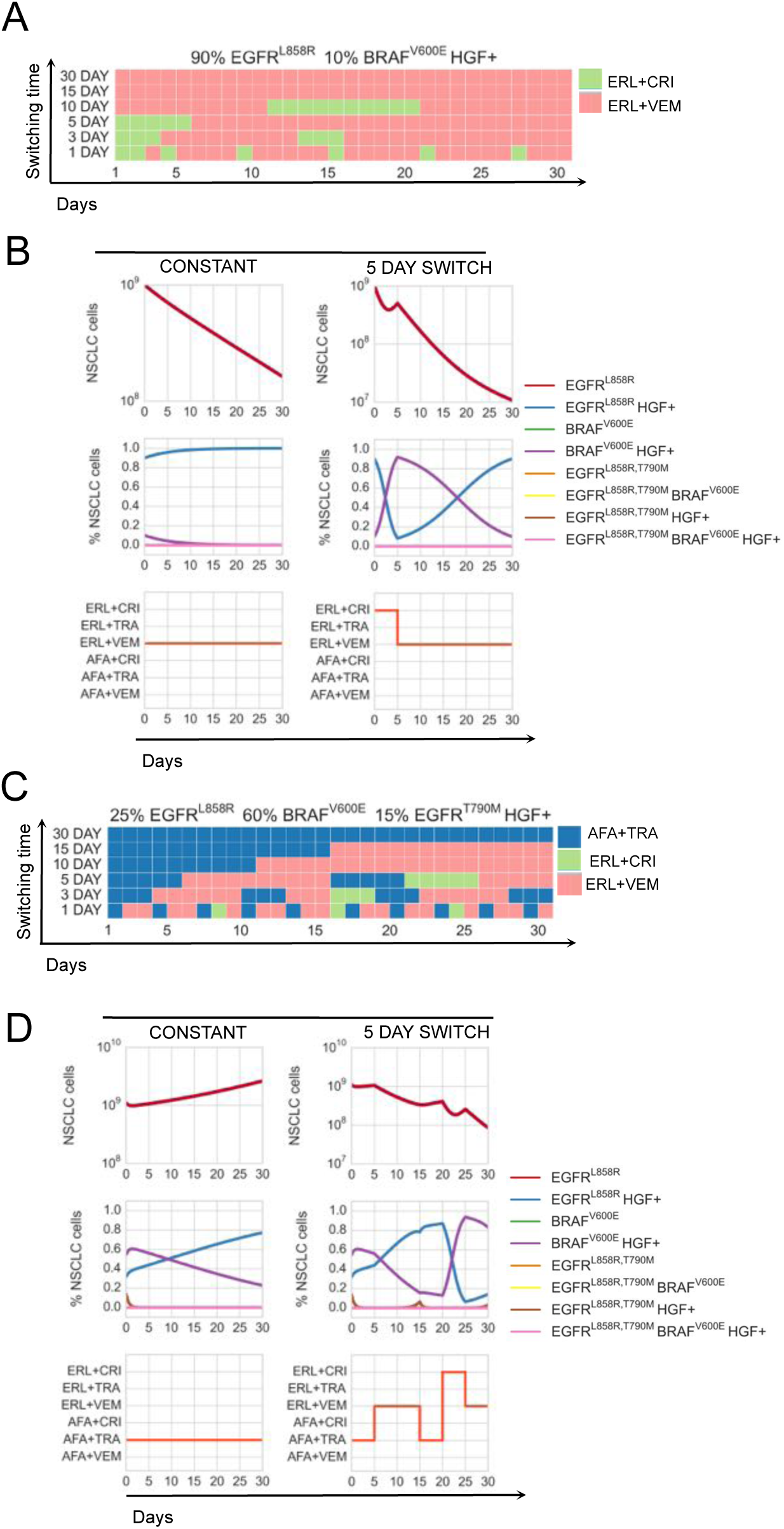
Optimal drug scheduling strategies solved by Algorithm 1 (SI, Section 2.2) for representative initial tumor cell distributions (A),(C), for a 30 day timeframe and 30, 15, 10, 5, 3 and 1 day minimum switching horizons, give one EGFR TKI, either 1.5 *µ*M erlotinib (ERL) or 0.5 *µ*M afatinib (AFA) in combination with either 5 *µ*M vemurafenib (VEM), 0.5 *µ*M trametinib (TRA) or 0.5 *µ*M crizotinib (CRI) and corresponding simulations (B),(D) of the lung adenocarcinoma evolutionary dynamics for a subset of optimal drug scheduling strategies.

**Table 1:**
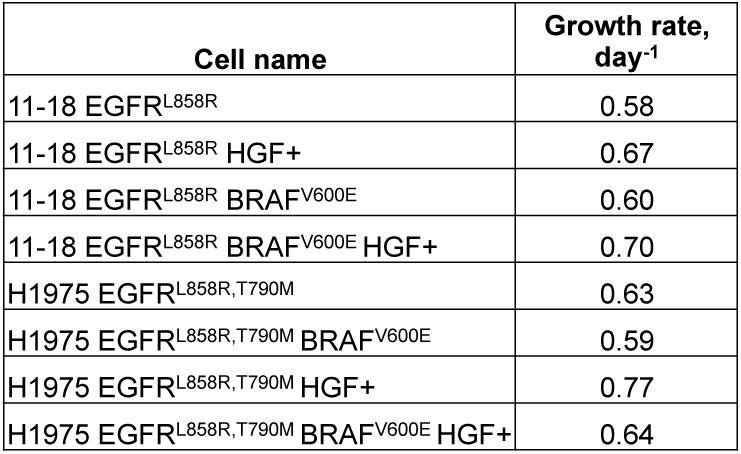
Experimentally derived growth rates in parental and engineered 11-18 EGFR^L858R^-positive lung adenocarcinoma cells and treated with or without HGF, fit with Equation (S8).

**Table 2:** Drug sensitivity as measured by the IC50 of erlotinib, afatinib, vemurafenib, trametinib and crizotinib in parental and engineered 11-18 EGFR^L858R^-positive lung adenocarcinoma cells.

| Cellname | IC50 in $\mu\text{M}$ | | | | |
| --- | --- | --- | --- | --- | --- |
|  | Erlotinib | Afatinib | Crizotinib | Trametinib | Vemurafenib |
| 11-18 EGFR<sup>L858R</sup> | 0.19 | 0.20 | 2.72 | 12.69 | 16.38 |
| 11-18 EGFR<sup>L858R</sup> HGF+ | 7.93 | 1.33 | 6.81 | 1.59 | 50.18 |
| 11-18 EGFR<sup>L858R</sup> BRAF<sup>V600E</sup> | 0.91 | 0.49 | 3.25 | 15.59 | 10.60 |
| 11-18 EGFR<sup>L858R</sup> BRAF<sup>V600E</sup> HGF+ | 8.74 | 1.49 | 10.54 | 1.49 | 12.64 |
| H1975 EGFR<sup>L858R,T790M</sup> | 7.54 | 0.08 | 9.33 | 0.76 | 48.31 |
| H1975 EGFR<sup>L858R,T790M</sup> BRAF<sup>V600E</sup> | 9.32 | 0.18 | 8.18 | 0.82 | 18.64 |
| H1975 EGFR<sup>L858R,T790M</sup> HGF+ | 7.04 | 0.60 | 25.59 | 0.12 | 53.89 |
| H1975 EGFR<sup>L858R,T790M</sup> BRAF<sup>V600E</sup> HGF+ | 7.97 | 0.82 | 31.83 | 0.06 | 54.48 |

**Table 3:** Drug sensitivity as measured by the IC50 of erlotinib in combination with 5 *µ*M vemurafenib, 0.5 *µ*M trametinib and 0.5 *µ*M crizotinib in parental and engineered 11-18 EGFR^L858R^-positive lung adenocarcinoma cells.

| Cell name | IC50 Erlotinib in $\mu\text{M}$ | | |
| --- | --- | --- | --- |
| | +0.5 $\mu\text{M}$ Crizotinib | +0.5 $\mu\text{M}$ Trametinib | +5 $\mu\text{M}$ Vemurafenib |
| 11-18 EGFR<sup>L858R</sup> | 0.30 | 0.19 | 0.30 |
| 11-18 EGFR<sup>L858R</sup> HGF+ | 0.18 | 0.47 | 1.34 |
| 11-18 EGFR<sup>L858R</sup> BRAF<sup>V600E</sup> | 1.64 | 0.08 | 0.09 |
| 11-18 EGFR<sup>L858R</sup> BRAF<sup>V600E</sup> HGF+ | 3.58 | 7.95 | 0.86 |
| H1975 EGFR<sup>L858R,T790M</sup> | 3.51 | 7.83 | 15.39 |
| H1975 EGFR<sup>L858R,T790M</sup> BRAF<sup>V600E</sup> | 3.71 | 7.68 | 9.86 |
| H1975 EGFR<sup>L858R,T790M</sup> HGF+ | 7.78 | 8.20 | 31.01 |
| H1975 EGFR<sup>L858R,T790M</sup> BRAF<sup>V600E</sup> HGF+ | 6.70 | 5.50 | 103.67 |

**Table 4:**
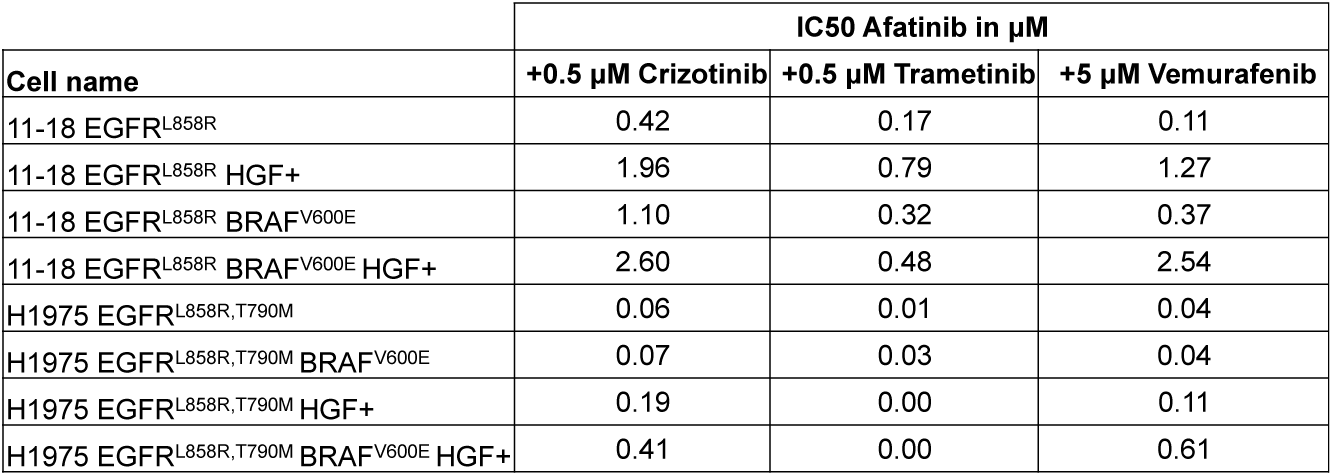
Drug sensitivity as measured by the IC50 of afatinib in combination with 5 *µ*M vemurafenib, 0.5 *µ*M trametinib and 0.5 *µ*M crizotinib in parental and engineered 11-18 EGFR^L858R^-positive lung adenocarcinoma cells.

| Cell name | IC50 Afatinib in $\mu\text{M}$ | | |
| --- | --- | --- | --- |
| | +0.5 $\mu\text{M}$ Crizotinib | +0.5 $\mu\text{M}$ Trametinib | +5 $\mu\text{M}$ Vemurafenib |
| 11-18 EGFR<sup>L858R</sup> | 0.42 | 0.17 | 0.11 |
| 11-18 EGFR<sup>L858R</sup> HGF+ | 1.96 | 0.79 | 1.27 |
| 11-18 EGFR<sup>L858R</sup> BRAF<sup>V600E</sup> | 1.10 | 0.32 | 0.37 |
| 11-18 EGFR<sup>L858R</sup> BRAF<sup>V600E</sup> HGF+ | 2.60 | 0.48 | 2.54 |
| H1975 EGFR<sup>L858R,T790M</sup> | 0.06 | 0.01 | 0.04 |
| H1975 EGFR<sup>L858R,T790M</sup> BRAF<sup>V600E</sup> | 0.07 | 0.03 | 0.04 |
| H1975 EGFR<sup>L858R,T790M</sup> HGF+ | 0.19 | 0.00 | 0.11 |
| H1975 EGFR<sup>L858R,T790M</sup> BRAF<sup>V600E</sup> HGF+ | 0.41 | 0.00 | 0.61 |

**Table 5:** Differential equation parameters derived using Equation (S11), corresponding to experimentally derived dose response curves of erlotinib and afatinib for parental and engineered 11-18 EGFR^L858R^-positive lung adenocarcinoma cells.

| Cell name | Erlotinib |  |  | Afatinib |  |  |
| --- | --- | --- | --- | --- | --- | --- |
| | $\gamma$ | n | K | $\gamma$ | n | K |
| 11-18 EGFR<sup>L858R</sup> | 2.59 | 0.54 | 1.22 | 73.45 | 0.32 | 424540.00 |
| 11-18 EGFR<sup>L858R</sup> HGF+ | 0.61 | 3.81 | 7.93 | 1885.80 | 0.70 | 106610.00 |
| 11-18 EGFR<sup>L858R</sup> BRAF<sup>V600E</sup> | 1.72 | 0.53 | 1.90 | 297.20 | 0.46 | 269500.00 |
| 11-18 EGFR<sup>L858R</sup> BRAF<sup>V600E</sup> HGF+ | 0.39 | 3.48 | 8.74 | 590.76 | 0.72 | 18112.00 |
| H1975 EGFR<sup>L858R, T790M</sup> | 1.87 | 2.23 | 9.54 | 3.77 | 1.49 | 0.22 |
| H1975 EGFR<sup>L858R, T790M</sup> BRAF<sup>V600E</sup> | 1.50 | 3.86 | 9.70 | 5.14 | 0.62 | 3.67 |
| H1975 EGFR<sup>L858R, T790M</sup> HGF+ | 0.52 | 3.67 | 7.04 | 250.96 | 0.56 | 21189.00 |
| H1975 EGFR<sup>L858R, T790M</sup> BRAF<sup>V600E</sup> HGF+ | 0.31 | 3.50 | 7.97 | 5.14 | 1.26 | 3.57 |

**Table 6:** Differential equation parameters derived using Equation (S11), corresponding to experimentally derived dose response curves of crizotinib, trametinib and vemurafenib for parental and engineered 11-18 EGFR^L858R^-positive lung adenocarcinoma cells.

| Cell name | Crizotinib |  |  | Trametinib |  |  | Vemurafenib |  |  |
| --- | --- | --- | --- | --- | --- | --- | --- | --- | --- |
| | $\gamma$ | n | K | gamma | n | K | $\gamma$ | n | K |
| 11-18 EGFR<sup>L858R</sup> | 8.80 | 1.84 | 10.36 | 207.05 | 0.94 | 5504.20 | 825.97 | 0.95 | 27377.00 |
| 11-18 EGFR<sup>L858R</sup> HGF+ | 6.14 | 3.89 | 11.57 | 3.47 | 1.29 | 4.67 | 9.81 | 4.42 | 89.86 |
| 11-18 EGFR<sup>L858R</sup> BRAF<sup>V600E</sup> | 74.60 | 1.59 | 61.34 | 1135.50 | 0.88 | 67225.00 | 672.41 | 0.69 | 236280.00 |
| 11-18 EGFR<sup>L858R</sup> BRAF<sup>V600E</sup> HGF+ | 235.65 | 0.57 | 270080.00 | 3.51 | 1.36 | 4.17 | 229.34 | 0.64 | 104240.00 |
| H1975 EGFR<sup>L858R, T790M</sup> | 6.98 | 1.68 | 34.62 | 28.40 | 0.26 | 1064100.00 | 1.56 | 5.88 | 50.20 |
| H1975 EGFR<sup>L858R, T790M</sup> BRAF<sup>V600E</sup> | 6.52 | 1.99 | 23.88 | 35.42 | 0.28 | 1223700.00 | 6.92 | 2.35 | 47.51 |
| H1975 EGFR<sup>L858R, T790M</sup> HGF+ | 4.66 | 2.51 | 51.31 | 47.17 | 0.27 | 900880.00 | 9.35 | 4.95 | 89.74 |
| H1975 EGFR<sup>L858R, T790M</sup> BRAF<sup>V600E</sup> HGF+ | 1.69 | 2.79 | 36.20 | 38.13 | 0.25 | 595280.00 | 8.62 | 4.92 | 89.37 |

**Table 7:** Differential equation parameters as derived using Equation (S11), corresponding to experimentally derived dose response curves of erlotinib in combination with either 0.5 *µ*M crizotinib, 0.5 *µ*M trametinib or 5 *µ*M vemurafenib for parental and engineered 11-18 EGFR^L858R^-positive lung adenocarcinoma cells.

| Cell name | Erlotinib+0.5 μM Crizotinib | | | Erlotinib+0.5 μM Trametinib | | | Erlotinib+5 μM Vemurafenib | | |
| --- | --- | --- | --- | --- | --- | --- | --- | --- | --- |
| | $\gamma$ | n | K | $\gamma$ | n | K | $\gamma$ | n | K |
| 11-18 EGFR<sup>L858R</sup> | 49.60 | 0.32 | 215970.00 | 23.91 | 0.38 | 2057.10 | 1.87 | 0.45 | 0.96 |
| 11-18 EGFR<sup>L858R</sup> HGF+ | 13.87 | 0.31 | 2167.30 | 197.39 | 0.44 | 180410.00 | 4.03 | 0.44 | 46.41 |
| 11-18 EGFR<sup>L858R</sup> BRAF<sup>V600E</sup> | 5.02 | 0.38 | 204.43 | 3.10 | 0.77 | 0.41 | 1.11 | 0.85 | 0.05 |
| 11-18 EGFR<sup>L858R</sup> BRAF<sup>V600E</sup> HGF+ | 3.79 | 0.75 | 26.45 | 1.28 | 0.95 | 6.69 | 1.97 | 0.95 | 1.63 |
| H1975 EGFR<sup>L858R, T790M</sup> | 990.17 | 0.74 | 65062.00 | 6.43 | 0.80 | 110.13 | 3.01 | 0.97 | 53.62 |
| H1975 EGFR<sup>L858R, T790M</sup> BRAF<sup>V600E</sup> | 3.40 | 1.22 | 11.30 | 2.29 | 4.29 | 9.33 | 2.20 | 1.54 | 16.34 |
| H1975 EGFR<sup>L858R, T790M</sup> HGF+ | 3.37 | 2.08 | 14.90 | 2.34 | 4.05 | 10.15 | 1.07 | 2.12 | 23.25 |
| H1975 EGFR<sup>L858R, T790M</sup> BRAF<sup>V600E</sup> HGF+ | 3.65 | 1.22 | 21.97 | 2.06 | 1.76 | 8.08 | 3.04 | 1.27 | 271.56 |

**Table 8:** Differential equation parameters derived using Equation (S11), corresponding to experimentally derived dose response curves of afatinib in combination with either 0.5 *µ*M crizotinib, 0.5 *µ*M trametinib or 5 *µ*M vemurafenib for parental and engineered 11-18 EGFR^L858R^-positive lung adenocarcinoma cells.

| Cell name | Afatinib+0.5 μM Crizotinib | | | Afatinib+0.5 μM Trametinib | | | Afatinib+5 μM Vemurafenib | | |
| --- | --- | --- | --- | --- | --- | --- | --- | --- | --- |
| | $\gamma$ | n | K | $\gamma$ | n | K | $\gamma$ | n | K |
| 11-18 EGFR<sup>L858R</sup> | 102.02 | 0.36 | 363520.00 | 190.37 | 0.42 | 116910.00 | 41.09 | 0.27 | 314410.00 |
| 11-18 EGFR<sup>L858R</sup> HGF+ | 212.01 | 0.49 | 209360.00 | 42.67 | 0.29 | 977910.00 | 66.90 | 0.36 | 344940.00 |
| 11-18 EGFR<sup>L858R</sup> BRAF<sup>V600E</sup> | 310.67 | 0.52 | 132530.00 | 141.93 | 0.60 | 2315.40 | 101.84 | 0.35 | 673450.00 |
| 11-18 EGFR<sup>L858R</sup> BRAF<sup>V600E</sup> HGF+ | 1440.80 | 0.69 | 161180.00 | 54.46 | 0.31 | 621930.00 | 311.18 | 0.61 | 57467.00 |
| H1975 EGFR<sup>L858R, T790M</sup> | 3.75 | 1.25 | 0.19 | 19.69 | 0.18 | 572790.00 | 3.87 | 0.77 | 0.26 |
| H1975 EGFR<sup>L858R, T790M</sup> BRAF<sup>V600E</sup> | 3.68 | 1.44 | 0.18 | 26.30 | 0.22 | 451080.00 | 3.70 | 1.07 | 0.17 |
| H1975 EGFR<sup>L858R, T790M</sup> HGF+ | 1549.80 | 0.77 | 4033.50 | 14.89 | 0.24 | 285.48 | 110.87 | 0.36 | 171470.00 |
| H1975 EGFR<sup>L858R, T790M</sup> BRAF<sup>V600E</sup> HGF+ | 4.59 | 1.88 | 1.02 | 35.76 | 0.23 | 34108.00 | 411.18 | 0.52 | 140300.00 |

